# Cryo-EM structures of human ClpXP reveal mechanisms of assembly and proteolytic activation

**DOI:** 10.1101/2024.11.12.623337

**Authors:** Wenqian Chen, Jie Yang, Gabriel C. Lander

## Abstract

The human ClpXP complex (hClpXP) plays a central role in mitochondrial protein quality control by degrading misfolded or unneeded proteins. While bacterial ClpXP complexes have been extensively characterized, the molecular determinants underlying hClpXP assembly and regulation are not as well understood. We determined cryo-electron microscopy (cryo-EM) structures of hClpP in isolation and in complex with hClpX, revealing how hClpX binding promotes rearrangement of an asymmetric hClpP heptamer to assemble as a symmetric tetradecamer. Our hClpXP structure also highlights the stabilizing role of a previously uncharacterized eukaryotic ClpX sequence, referred to as the E-loop, and its importance in ATPase activity and hexamer assembly. We further show that peptide interaction with the hClpP proteolytic active site promotes the complex to adopt a proteolytically competent conformation. Together, these findings advance our understanding of the molecular mechanisms defining hClpXP activation and function.

## Introduction

Mitochondria are busy signaling hubs where a multitude of cellular processes converge, including oxidative phosphorylation, calcium homeostasis, redox signaling, and apoptosis^1–3^. Maintaining internal protein homeostasis (proteostasis) involves a sophisticated and tightly controlled protein quality control network that ensures proper import, folding, and turnover of proteins^4^. In humans, disruptions in this network are associated with aging and can lead to a wide range of diseases, including cancer, neurological disorders, diabetes, and skin disorders^5^. The ClpXP complex is one of the key ATP-dependent proteases involved in this protein quality control network, specifically targeting and degrading misfolded, damaged, or surplus proteins in the matrix of mitochondria. While all other mitochondrial proteases are encoded by a single gene, ClpXP is an assembly of two distinct proteins: ClpX, an ATPase Associated with diverse cellular activities (AAA+) with unfoldase activity, and ClpP, a conserved serine protease. Together, these components work in concert to effectively degrade damaged proteins, as well as regulating essential cellular functions, including heme biosynthesis^6,7^, mitochondrial DNA nucleoid distribution^8^, and reactive oxygen species (ROS) production^9^.

Despite extensive biochemical and structural characterization of unicellular ClpXP systems^10–14^, many of the molecular details underlying human ClpXP (hClpXP) assembly and proteolytic activation are uncharacterized. ClpXP systems are broadly conserved across all forms of life, but the specific requirements of eukaryotic proteostasis have driven ClpX to the evolve distinct regulatory components. For example, hClpX features a unique eukaryotic-specific insertion within its ATPase domain, previously termed the E-domain^15^ (**Supplementary Fig. 1a**). The function of this insertion was unknown, but its positioning at the distal end of hClpX suggests a potential function in substrate recognition^16^ (**Supplementary Fig. 1c**). Additionally, hClpP possesses an additional 28 amino acids at its C-terminus, termed the C-terminal extension (**Supplementary Fig. 2b**). Deleting this extension promotes tighter interaction between hClpP to hClpX^17^, but the precise underlying mechanism remains unclear.

The ClpP protease has also diverged through evolution. Bacterial ClpP assembles as a tetradecamer composed of two stacked heptameric rings that enclose 14 proteolytic active sites within the chamber^18^, but this is not necessarily the case for hClpP. Some studies suggest that hClpP exists as a tetradecamer^19–21^, whereas others indicate that ClpP assembles as a stable heptamer and only forms a tetradecamer upon binding with hClpX^17,22^. Studies employing small molecule activators of hClpP have provided some insights into its mechanism of activation^21,23,24^, but in the absence of a direct structural comparison between apo hClpP and hClpX-bound hClpP, the precise mechanism of hClpP activation remains elusive.

To gain insights into the molecular mechanisms that regulate hClpXP function, we determined the cryo-EM structures of hClpP on its own and in complex with hClpX. Our hClpXP structures revealed the architecture of the E-domain (hereafter referred to as the E-loop), which we show to be involved in nucleotide sensing and hClpX hexamer stabilization. Importantly, our structural analyses reveal that, unlike its bacterial homologs, hClpP assembles as an asymmetric heptamer and undergoes a conformational rearrangement to accommodate symmetric tetradecamer formation upon hClpX binding. After introducing a peptide mimetic to hClpP, we demonstrate that substrate interaction with the proteolytic active site drives a rearrangement that positions the catalytic residues for peptide hydrolysis. Finally, we show that the C-terminal extension of hClpP negatively regulates the interaction between hClpX and hClpP by physically obstructing the hydrophobic pockets on hClpP where hClpX binds. Together, these findings highlight the allosteric mechanisms that have evolved for hClpXP to meet the complex demands of the mitochondrial environment.

## Results

### Cryo-EM structure of hClpXP reveals a conserved substrate translocation mechanism by hClpX

We first aimed to compare the structure of the hClpXP complex to the numerous prior structures of the unicellular homologs to identify evolutionarily conserved or divergent features. Assembling the complex for structural studies required extensive sample optimization (see methods), but we ultimately succeeded in determining the structure of the hClpXP complex at an overall resolution of ~3 Å (**Fig. 1a, Supplementary Figs. 3–4**, and **Table 1**). The reconstruction is generally reminiscent of previously determined bacterial ClpXP structures, including a co-purified peptide bound within the central pore of the hClpX unfoldase, a common feature of recombinantly expressed AAA+ proteins containing a Walker B mutation^12,25,26^. The hexameric assembly of hClpX adopts a shallow spiral conformation consistent with bacterial ClpX and other AAA+ unfoldases^11,12,14^, and we labeled the subunits of hClpX in alphabetical order (A, B, C, D, E, and F) with chain A at the uppermost position of the spiral and continuing downwards, and chain F identified as the displaced “seam” subunit disengaged from the peptide substrate **(Fig. 1b**). Also consistent with other AAA+ unfoldases was density corresponding to an ATP nucleotide with a coordinated magnesium cofactor observed in the binding pockets of chains A through D, where the ATP β- and γ-phosphates are stabilized by interactions with K299 (Walker A motif) and R562 (sensor-II) of the same subunit, as well as R499 (arginine finger) from the adjacent protomer (**Supplementary Fig. 5**). The nucleotide density in lowest subunit of the spiral, chain E, was representative of ADP and lacked observable magnesium density (**Supplementary Fig. 5b**). Further, the arginine finger from the neighboring F subunit is shifted away to produce a more open nucleotide-binding pocket. No nucleotide density was resolved in the seam subunit F.

**Table 1.**
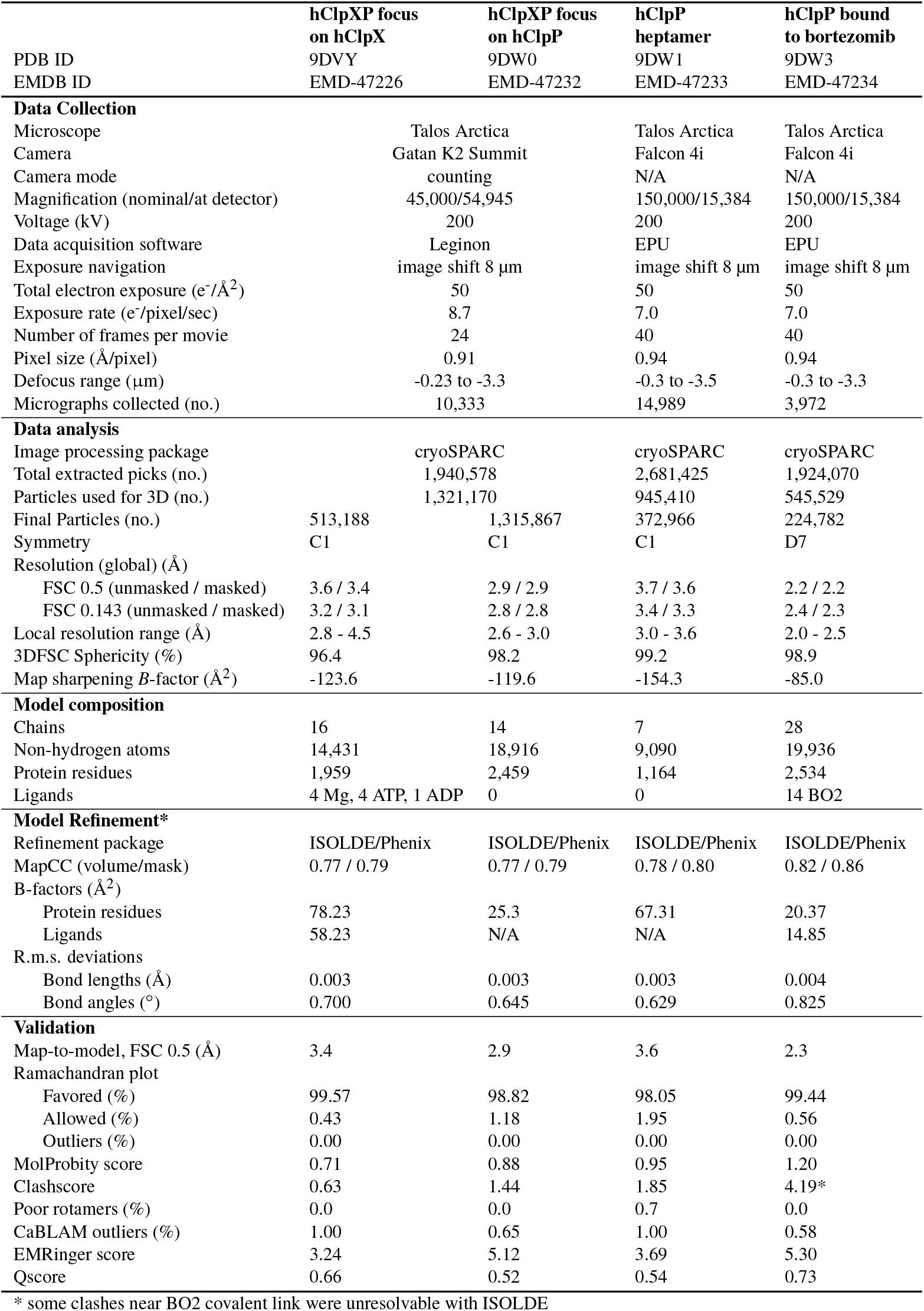
Cryo-EM data collection, refinement, and modeling statistics for hClpX, hClpX-bound-hClpP, hClpP heptamer, and bortezomib-bound hClpP.

**Figure 1.**
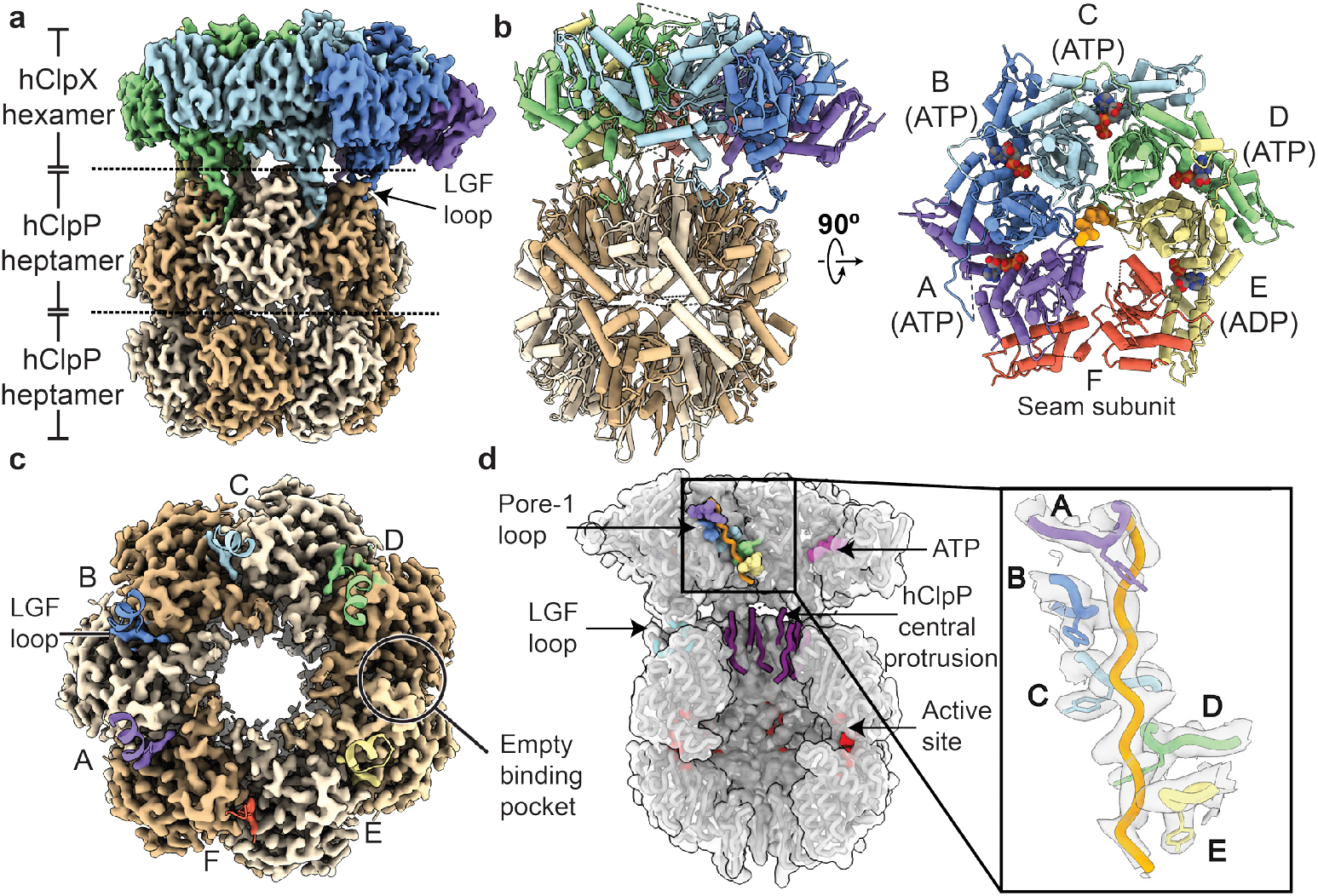
Cryo-EM reconstruction of human mitochondrial ClpXP. (**a**) Cryo-EM map of the hClpXP complex with a substrate peptide bound. Each subunit of the hClpX homo-hexamer is colored distinctly. (**b**) Atomic structural model of hClpXP. The substrate peptide within the hClpX central pore is modeled as poly-alanine. The hClpX subunits are labeled A-F with the nucleotide states indicated. Chain F, the seam subunit, is disengaged from the substrate. (**c**) Top view of the hClpP map with the hClpX LGF loops (shown as helices) occupying the hydrophobic pocket. An empty LGF loop binding pocket is located between chains D and E. (**d**) Cut-away view of the hClpXP 3D reconstruction highlighting several features: pore-1 loops, nucleotides, LGF loops, the hClpP central protrusion, and the active site. The inset shows Y327 from the pore-1 loops forming a spiral staircase around the substrate peptide backbone.

As previously observed in other AAA+ proteins, the nucleotide state of each protomer correlates with the extent of the subunit’s interaction with the incoming substrate within the central ATPase channel^27–29^. The pore-1 loop of each nucleotide-bound hClpX subunit in the right-handed spiral engages the translocating peptide via a conserved GYVG sequence, so that the aromatic-hydrophobic motif intercalates into the substrate backbone (**Fig. 1d**). This organization maintains grip on the incoming polypeptide to facilitate translocation towards the peptidase chamber using a putative hand-over-hand mechanism^11,12,28–30^.

ClpX attaches to ClpP via loops containing a conserved Ile/Leu-Gly-Phe (IGF or LGF) motifs, which bind to the hydrophobic pockets formed at the inter-protomer ClpP interface^11^. Our hClpXP structure shows how F441 of the hClpX LGF loop forms conserved hydrophobic interactions with residues Y118, W146, Y138, and L104 in the hydrophobic pocket of hClpP (**Supplementary Fig. 6**). Prior cryo-EM studies of bacterial ClpXP systems have shown that one of the hydrophobic binding pockets in ClpP is empty due to the six-to-seven protomer mismatch between the unfoldase and protease. However, this empty pocket is sometimes situated between the clefts occupied by the IGF loops of ClpX subunits E and F^11,12,14^ while in others it is located between subunits D and E^31,32^. It was proposed that the ATP hydrolysis-driven motions of the ATPase subunits coordinate repositioning of the ClpX I/LGF loops as well as the stability of the N-terminal β-hairpins comprising the ClpP axial channel to facilitate substrate translocation^11^. The pore-2 loops of ClpX are also thought to coordinate substrate translocation through interactions with the axial ClpP β-hairpins that vary based on ClpX nucleotide state^11-13,33^. In our hClpXP reconstruction, we observe an empty pocket between hClpX chains D and E (**Fig. 1c)**, and although our pore-2 loops are not resolved, we observe that the pore-2-adjacent residue D374 from hClpX subunit E interacts with R68 at the top of a hClpP β-hairpin (**Supplementary Fig. 7**). This observation is similar to the *Neisseria meningitides* ClpXP structure, where only a single ClpX protomer (subunit E) interacts with the arginine residue from ClpP β-hairpin at the same position^11^, suggesting a conserved ClpXP communication and substrate translocation mechanism.

### E-domain of hClpX senses nucleotide states and promotes hexamer formation

A notable distinguishing feature of our hClpXP structure is the presence of the E-loop, a eukaryotic-specific insertion of unknown function within the AAA+ module (**Fig. 2a**). Within the hClpX hexamer, the E-loop extends from the large AAA+ domain of one subunit over the adjacent nucleotide binding pocket and interacts with the neighboring subunit at the interface of the large and small AAA+ domains. Despite these numerous interactions, the E-loop is generally flexible, and is only partially resolved. The interface between the large and small AAA+ domains is distinct for each subunit within translocating AAA+ unfoldases due to the staircase organization of the hexamer^27–29^, thereby impacting E-loop interactions. Accordingly, the resolvability of the E-loop density varies among different subunits, indicating that the register of the subunit within the hexamer impacts the E-loop stability (**Supplementary Fig. 8b**). The least-resolved E-loop is that of the seam subunit, chain F, which extends over the ADP-bound pocket of chain E. Proceeding from the top of the ATPase spiral to the bottom, the E-loop arching across each of the preceding nucleotide binding pockets becomes increasingly ordered, with the E-loop extending over the lower-most ATP pocket being the best-resolved of the loops. Given the location of the E-loop, and that E285 within the loop is within hydrogen bonding distance of both the nucleotide ribose and R565 in the binding pocket (**Fig. 2a** and **Supplementary Fig. 8a**,**b**), we surmised that the E-loop may play a role in the ATPase mechanochemical cycle.

**Figure 2.**
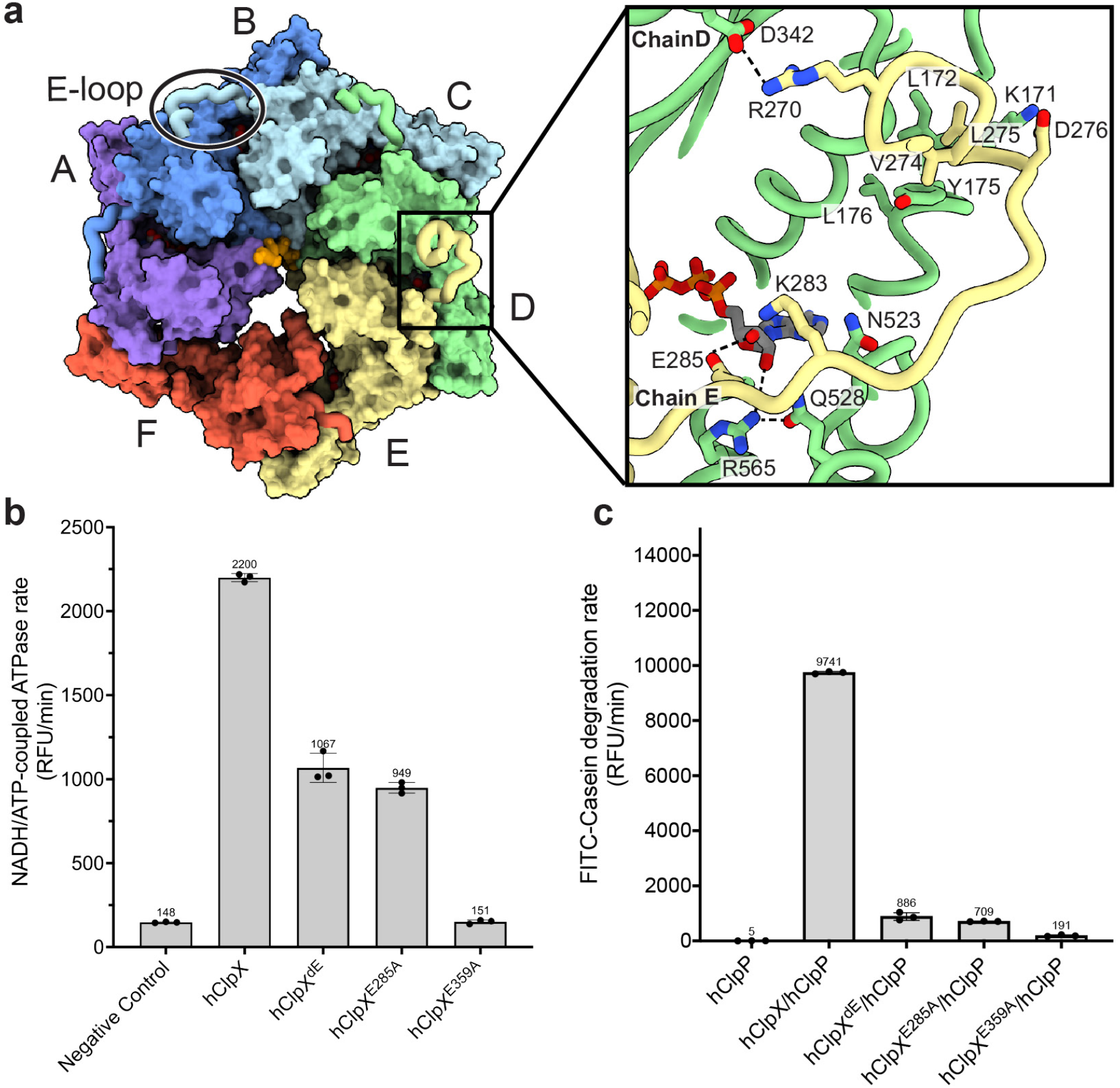
The hClpX E-loop senses nucleotide states and facilitates inter-subunit communication. (**a**) The hClpX hexamer is shown as a surface representation, highlighting the E-loop (shown as licorice/ovals style) of one hClpX subunit extends away to interact with the neighboring subunit. The inset illustrates the network of interactions between the E-loop of chain E and chain D, emphasizing that E285 from chain E is within hydrogen bonding distance of the ATP nucleotide. (**b**) NADH/ATP-coupled ATPase assays were performed. The negative controls include reactions without enzymes and the Walker B mutant—hClpX^E395A^. Removal of the E-loop from hClpX caused a 2-fold decrease in hClpX ATPase activity. The E285A mutation has similar deleterious effect on hClpX ATPase activity as the E-loop deletion. Data are presented as bar graphs showing mean values with error bars representing standard deviation from three independent replicates. (**c**) Removal of the E-loop from hClpX or introduction of the E285A mutation resulted in a 10X reduction in the FITC-casein degradation activity of the hClpXP complex. Data are presented as bar graphs showing mean values with error bars showing standard deviation from three independent replicates.

To test this hypothesis, we removed the E-loop (residues 230-304) from hClpX (hClpX^dE^) and evaluated its impact on ATPase activity and substrate degradation. Deletion of the E-loop decreased ATPase activity by more than half and impaired the ability of hClpXP to degrade casein substrates, as indicated by a 11-fold reduction in FITC-casein degradation activity (**Fig. 2b,c**). Negative stain EM analysis of this truncation mutant revealed an inability of the hClpX^dE^ to form hexamers, instead assembling into higher-order oligomers, including heptameric, octameric, and nonameric assemblies in the absence of exogenous ATP (**Supplementary Fig. 8d**). Although the addition of ATP and MgCl_2_ facilitated the formation of some loosely organized hexamers, we also observed heptamers and unclosed ring assemblies. Notably, site-specifically mutating the nucleotide-interacting residue E285 within the E-loop to alanine was as deleterious to hClpXP enzymatic activities as removal of the E-loop, but the mutation did not negatively affect hClpX hexamer formation (**Fig. 2b,c** and **Supplementary Fig. 8e**). These results suggest that the E285 residue plays a key role in hClpX ATPase and substrate translocation activities by sensing and communicating changes in nucleotide states between neighboring subunits.

Similar to unicellular ClpX systems, ATP binding is important for hClpX to form hexamers that are competent for substrate translocation^16^ (**Supplementary Fig. 8c**). However, our data indicate that the E-loop serves as a critical structural feature required for proper hexameric assembly of hClpX and effective ATP-driven substrate translocation. Given that the E-loop is specific to eukaryotes, these findings suggest an evolutionary adaptation of hClpX to incorporate a more complex set of structural requirements for stabilizing hexamer formation and sensing nucleotide states during substrate translocation in mitochondria. This is consistent with our observation that the hClpX complex does not assemble into hexamers or the hClpXP holo-complex as readily as its bacterial homologs (**Supplementary Fig. 3**).

### hClpP assembles as an asymmetric heptamer whose C-terminal extension negatively regulates its activity

Having probed the evolutionarily distinct features of the assembled hClpXP complex, we next aimed to investigate the mechanism of hClpXP assembly. It is well-established that bacterial ClpP proteins form stable tetradecamers composed of two stacked heptamers, but there are conflicting results regarding whether hClpP assembles as heptamer or tetradecamer in the absence of hClpX in solution^17,20-22^. When we expressed and purified our wild type hClpP construct, we observed that it eluted from size exclusion chromatography at a mass corresponding to a heptameric assembly (**Supplementary Fig. 9a**). We further confirmed that hClpP predominantly exists as a heptamer in solution using mass photometry and negative-stain EM (**Supplementary Fig. 9b-c**), and next aimed to determine a high-resolution cryo-EM structure. We observed that the apo hClpP heptamer exhibits a strong preferred axial orientation when vitrified on cryo-EM grids (**Supplementary Fig. 10a**,**b**), which was alleviated by the addition of fluorinated fos-choline-8 detergent (**Supplementary Fig. 10c**). Although the majority of imaged hClpP oligomers were heptameric according to 2D classification, there was a minor population of tetradecameric hClpP, possibly induced by the high concentration of hClpP applied to the cryoEM grid (15 mg/ml). Our 2D averages also showed that the tetradecamers were not well-resolved (circled in red in **Supplementary Fig. 10d**), indicative of flexibility of transient inter-heptamer interactions. These data indicate that the hClpP heptamers do not readily form stable tetradecamers, suggesting a role for hClpX in promoting assembly of the hClpP tetradecamer.

Image analysis of the hClpP heptamer was challenged by the reduced apparent particle concentration and increased ice thickness associated with introducing detergent to the sample, but we obtained a heptameric hClpP reconstruction with a reported overall resolution of ~3.5 Å (**Supplementary Fig. 10d**,**e** and **Table 1**), which was sufficient to generate an atomic model (**Fig. 3a**,**b**). Surprisingly, we found that the hClpP heptamer is not symmetric as expected, but rather assembles into an asymmetric oligomer in the absence of hClpX (**Supplementary Fig. 11**). A salient feature of the asymmetric heptamer is the ordering of three of the N-terminal loops into β-hairpins, forming a partial wall around the axial hClpP pore, while these loops from the other four subunits are disordered (**Fig. 3b**). We surmise that this distinct asymmetric organization of the N-terminal loops forming a partial wall at the ClpP pore impacts hClpX recruitment, although further biophysical studies will be required to elucidate the possible regulatory mechanism of this unexpected structural feature.

**Figure 3.**
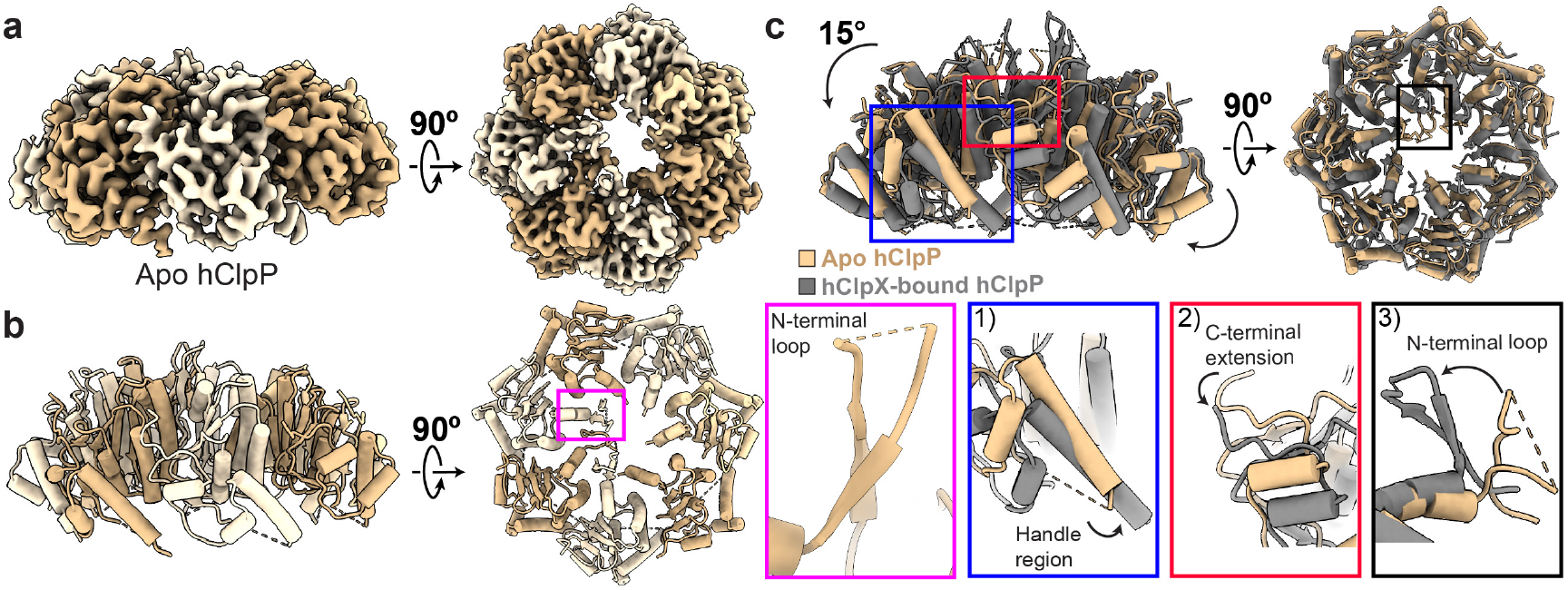
Apo hClpP undergoes key conformational changes upon binding to hClpX. (**a**) 3D reconstruction and (**b**) atomic model of C1 refined apo hClpP. The N-terminal loops of three hClpP subunits form well-ordered β-hairpins and are pointed upwards, while the N-terminal loops of the remaining subunits are disordered. The inset highlighted with a magenta box shows one of the three well-ordered N-terminal β-hairpins (**c**) Overlay of the apo hClpP (shown in burlywood) and the top part of the hClpX-bound hClpP tetradecamer (shown in grey). The insets highlight conformational changes in three different regions of hClpP upon binding to hClpX: 1) inward movement of the handle region; 2)a downward movement of the C-terminal extension; 3) outward movement of the axial loops from the central pore as well as the ordering of the N-terminal loops in the subunits that originally had disordered loops.

hClpP contains an unstructured 28-residue C-terminal extension (**Supplementary Fig. 2b**) that is unresolved in our hClpP reconstructions. We wondered how the removal of this disordered region from ClpP increases affinity for ClpX and increases substrate degradation^17^ (**Supplementary Figs. 3b, 12c**). Prior studies of the C-terminal extension in ClpP homologs from *Mycobacterium smegmatis*^34^ and *Listeria monocyotenes*^13^ suggest that this segment may be positioned to block the interaction between the ClpX loops and their respective hydrophobic binding pockets in ClpP. In our apo hClpP structure, we noted that the two proline residues preceding the C-terminal extension are well-resolved, stably positioned atop the LGF loop binding pocket. The location of these proline residues confirms that the C-terminal extensions are located in the vicinity of the hClpP LGF loop binding pockets, where they could potentially shield the pockets from hClpX interaction. Further, these prolines residues are positioned so that they would sterically hinder the binding of LGF loops (**Supplementary Fig. 12a**). We noted that these residues are shifted away from the LGF binding pocket in our hClpXP structure to accommodate the interactions necessary to form the heterocomplex (**Supplementary Fig. 12b**). These structural observations support a conserved mechanism wherein the C-terminal extension negatively regulates the interaction between hClpX and hClpP, requiring its displacement to accommodate hClpX binding. This regulatory function of the C-terminal extension may be necessary for the hClpP-independent function of hClpX as a chaperone^6,7^, perhaps preventing non-specific degradation of substrates.

### hClpP switches from an asymmetric heptamer to a symmetric tetradecamer upon interaction with hClpX

In addition to displacing the C-terminal extension of hClpP, hClpX binding is associated with a large-scale reorganization of the hClpP heptamer. Superimposing our structures of the apo hClpP heptamer and one heptameric ring of the hClpX-bound hClpP reveals several notable conformational changes (**Fig. 3c**). Upon hClpX binding, the four N-terminal loops that were disordered in our asymmetric heptameric hClpP structure adopt the canonical β-hairpin fold, concurrent with expansion of the axial pore to allow unfolded substrates to access the proteolytic chamber.

Notably, we also observe a ~15° axial rotation of the subunits away from the hClpX hexamer as the hClpP heptamer transitions from an asymmetric heptamer to a symmetric assembly (**Fig. 3c**). The rotation of the hClpP subunits upon hClpX binding results in an inward movement of the handle regions to facilitate critical interactions at the hClpP-hClpP interface, promoting a stable tetradcamer conformation (**Supplementary Fig. 14a**). In the hClpX-bound hClpP tetradacameric structure, the conserved oligomerization sensors E225 and R226 from one ClpP heptamer are positioned in proximity to form salt bridges with R226 and E225 from the opposing ClpP heptamer (**Supplementary Fig. 14a**). When we introduced the E225A and R226A mutation to hClpP, these mutations destabilize, but do not entirely abolish, hClpP tetradcamer formation in the presence of hClpX, as assessed by negative stain EM (**Supplementary Fig. 14c**). We also observe several other residues in the handle region, including E196, K202, and Y206, that interact with residues from the opposing subunit to maintain the tetradacameric state of hClpP. Additionally, the catalytic residue D227 contributes to stabilizing the tetradecamer by forming a salt bridge with Q194. These interactions were also recently observed in a tetradecameric cryoEM structure of hClpP determined while this manuscript was under preparation^35^.

### Substrate binding is required for hClpP activation

Prior structural studies of bacterial ClpP revealed three distinct states of the tetradecameric assembly, defined as extended, compact, and compressed^18^. The extended state is considered to be the proteolytically active conformation, whereas the compact and compressed states are regarded as transition states posited to facilitate peptide exit^10,18^. However, in seeming contradiction to these results from unicellular systems, the crystal structure of apo hClpP revealed a tetradecamer in an extended state^19^, while crystal structures of hClpP bound to small molecule activators showed tetradecamers in a compact state^21,23,24,36^. While it is possible that hClpP has evolved a fundamentally distinct proteolytic mechanism, it is also conceivable that these structural conformations were not representative of their corresponding proteolytic states due to crystallization conditions. We thus aimed to clarify these structure-activity relationships with cryo-EM analyses. Prior single particle cryo-EM studies of bacterial ClpXP complexes indicate that ClpX binding promotes ClpP to adopt the proteolytically competent extended conformation^10^. Given the level of structural conservation observed between bacterial ClpXP and our hClpXP complexes, we expected this feature to be preserved. Intriguingly, our hClpXP complex, despite being poised for degradation with a substrate peptide in the hClpX central pore, contains hClpP in a compact conformation that is presumed to be proteolytically inactive (**Fig. 4a**). In this inactive conformation, the catalytic histidine is orientated away from the nucleophilic serine such that the necessary hydrogen bond network required for catalysis cannot be achieved^37^.

**Figure 4.**
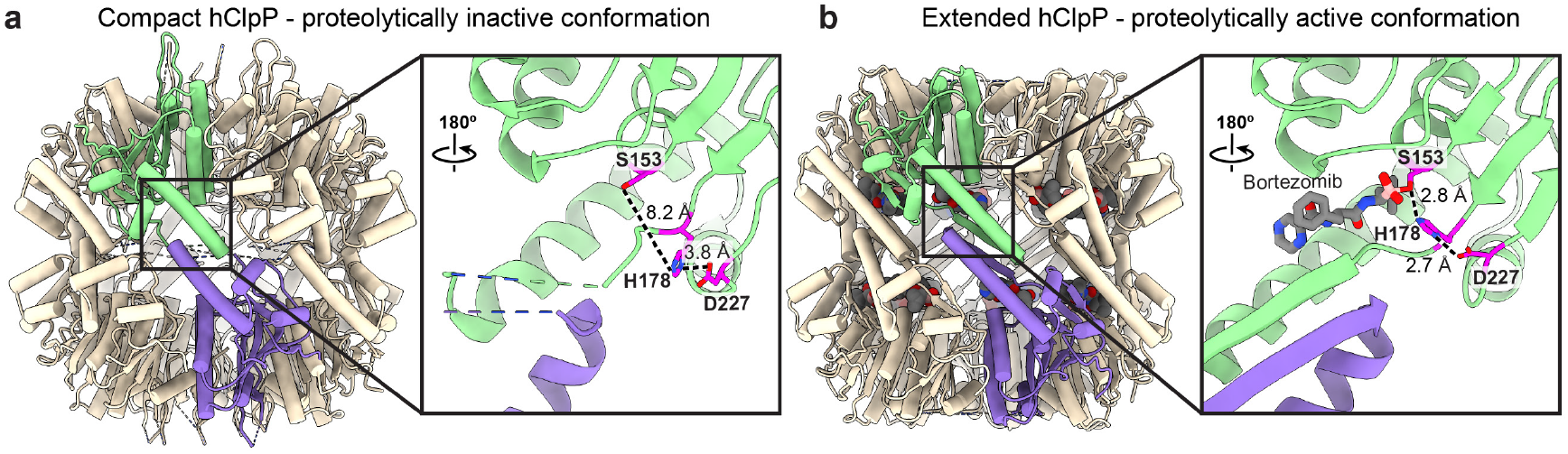
Bortezomib binding promotes hClpXP to adopt a proteolytically active conformation. (**a**) Atomic model of the compact hClpX-bound hClpP (**b**) Atomic model of the extended bortezomib-bound hClpP tetradecamers, with insets highlighting the active site conformations. To determine our hClpX-bound hClpP structure, S153 was mutated to alanine, but is shown here as a serine for visualization purposes. In this proteolytically inactive conformation, the imidazole of H178 is positioned too far from S153 for peptide hydrolysis. Conversely, in the bortezomib-bound hClpP, the active site adopts a proteolytically active conformation, with H178 shifting towards S153 in proximity for catalysis.

We previously observed an analogous discrepancy between proteolytically competent conformations in our comparison of the bacterial and human Lon protease^27^. Substrate interaction with the AAA+ module of bacterial Lon promotes rearrangement of the protease component to an active conformation, but insufficient for protease activation in human LONP1. We previously showed that the proteolytically active conformation of LONP1 could be obtained by introducing the peptide mimetic bortezomib^27^, and we surmised that hClpP activation may similarly require substrate peptide interaction with the peptidase to promote the active/extended conformation. We next determined a cryo-EM structure of bortezomib-bound hClpP to an overall resolution of ~2.3 Å, which was sufficient to resolve the covalent linkage between bortezomib and the catalytic S153 residue at the protease site (**Fig. 4b** and **Supplementary Fig. 13**). Notably, bortezomib binding induced hClpP to adopt an active, extended tetradecamer, confirming our hypothesis.

The similarity between our bortezomib-bound hClpP structure and the previous X-ray crystal structure of apo hClpP may be explained by the presence of an unknown density linked to the catalytic serine residue in the crystal structure, which may have contributed to adoption of the active conformation^19^. The bortezomib density in our structure also resembles that of a putative tripeptide identified in the protease active site of *Thermus thermophilus* ClpP, which also crystallized in the active conformation^38^. These structural data help clarify the interpretation of these prior crystallographic studies, and collectively demonstrate a conserved role for substrate peptide interaction with the proteolytic active site in protease activation. The conformational impact of substrate interaction is also consistent with findings from another cryo-EM study of hClpP that was performed while this manuscript was under preparation^35^.

## Discussion

ClpXP is a conserved protease that is critical for proteostasis in organisms belonging to all the kingdoms of life. While previous studies have shed light on the mechanisms that regulate substrate processing in bacterial ClpXP, it was unclear if human ClpXP employed these same mechanisms in the mitochondria. The specific proteostatic demands of the mitochondrial environment are distinct from those of unicellular organisms, and our findings reveal both conserved and evolutionarily distinct regulatory features that govern hClpXP function. While hClpXP shares a similar mechanism of substrate unfolding and degradation to its bacterial homologs, the assembly and activation of this mitochondrial protease are unique, involving a multi-tiered regulatory mechanism. The complexity of hClpXP regulation has likely evolved as a means of rapidly responding to the constantly fluctuating demands of mitochondria required to ensure overall cellular health.

Among the most striking of our observations is that, unlike previously characterized ClpP homologs that assemble as a symmetric tetradecamer, hClpP assembles as an asymmetric heptamer. In bacteria, the ClpP protease sites are sequestered within the tetradecameric chamber of ClpP, where flexible N-terminal loops restrict substrate access until ClpX binds, whereupon N-terminal gates of ClpP allow unfolded substrate peptides to enter and be degraded^39,40^. In contrast, the proteolytic active sites of the hClpP heptamer are exposed to the mitochondrial matrix. This heptameric assembly represents the first layer of regulation in the multi-tier regulatory system for hClpXP-mediated substrate degradation (**Fig 5**). The catalytic residues of the protease are positioned such that they are incompatible for proteolysis, preventing non-specific degradation of proteins. Both ClpX-induced conformational rearrangements, as well as interactions with another heptamer, are necessary to achieve proteolytic competence. This sophisticated mechanism ensures that protease activity is tightly coupled to cofactors such as ClpX, which are themselves likely to be tightly regulated.

**Figure 5.**
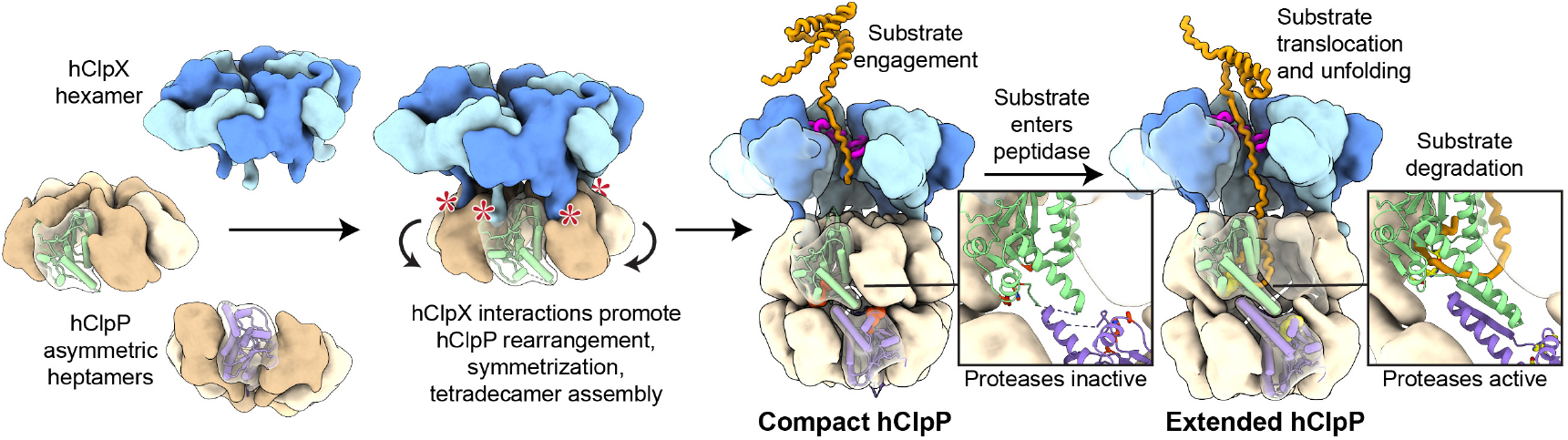
hClpXP assembly and activation mechanism. Cartoon summary of hClpXP activation mechanism. Apo hClpP predominantly exists as an asymmetric heptamer. Binding of hClpX and the substrate induces the formation of a symmetric, compact hClpP tetradecamer. Activation of hClpP occurs only when the substrate binds to the active site.

One such potential regulatory mechanism may involve hClpX assembly. We showed that the assembly of hClpP into a functional tetradecamer is promoted by interaction with the hClpX hexamer. However, hClpX hexamerization in vitro required substantial construct and condition optimization, suggesting that hClpX hexamerization is unlikely to be spontaneous in the mitochondrial matrix and likely only occurring in response to specific stimuli. The hClpX NTD contains a conserved C4-type zinc-finger motif known to be involved in adaptor binding and substrate recruitment^16,41^. Removing the NTD from hClpX enhanced its stability and facilitated hexamer formation, suggesting that the NTD may inhibit hClpX oligomerization unless it is bound to a mitochondrial adaptor protein or specific substrate. For example, interaction between the hClpX NTD and adaptor protein polymerase delta-interacting protein (PDIP38) was shown to limit hClpX turnover by LONP1 in mitochondria^41^. PDIP38 may interact with the hClpX NTD to promote hexamerization and ClpXP assembly, explaining the stabilizing role this adaptor protein has on hClpX level^41^.

The next layer of regulation involves the interaction between hClpX and the hClpP, which also required construct design and crosslinking to stabilize in vitro, suggesting that specific conditions are required for hClpXP complex assembly in the mitochondrion. Similar to hClpX hexamerization, hClpXP formation may also have substrate or cofactor requirements to ensure degradation of appropriate substrates. The more transient nature of the hClpX-hClpP interaction compared to bacterial homologs may be associated with the role hClpX plays as a chaperone in the mitochondrial matrix. For example, hClpX activates ALA synthase (ALAS) by partially unfolding the protein to facilitate binding of a cofactor heme^6^, yet under heme-repleting conditions, ALAS becomes targeted for hClpXP-mediated degradation^42^. To balance these dual chaperoning and unfoldase roles, regulatory checkpoints such as displacement of the hClpP C-terminal extension have likely evolved to enable substrate sensing and molecular decisions regarding unfolding, chaperoning, and degradation to be carried out effectively.

Once fully assembled, the hClpXP complex bears a conserved organization to unicellular ClpXP structures that likely operates through a conserved mechanochemical cycle. However, the hClpX AAA+ domain has integrated an E-loop that is critical for stabilizing the hexamer and carrying out the ATP hydrolysis associated with substrate unfolding and translocation. Unique motifs or insertions that facilitate inter-subunit communication or nucleotide sensing are found throughout the different classes of AAA+ modules. For example, AAA+ enzymes of the classical clade, such as YME1 and AFG3L2, harbor an inter-subunit signaling (ISS) motif that detects the nucleotide state of the adjacent subunit^25,28^. ClpX is a member of the HCLR clade, which all contain a presensor-1 β-hairpin (PS1βH) that has been shown in other members of the clade to be crucial for AAA+ function such as sensing nucleotide states^29^ or stabilizing the pore-1 loop during substrate translocation^27^. However, the PS1βH sequence is highly divergent (**Supplementary Fig. 2a**) and seems to play varying functional roles AAA+ members of the PS1βH superclade^43^. It is plausible that the nucleotide sensing role of PS1βH in hClpX has been evolutionarily transferred to the E-loop. Additionally, this addition to the hClpX AAA+ module may help fine-tune the rate of ATP hydrolysis or inter-subunit coordination in response to different substrates under varying mitochondrial conditions^26^.

Finally, in contrast to bacterial systems, hClpX binding is insufficient to induce the active conformation of hClpP. While hClpX binding drives hClpP to form compact, symmetric tetradecamers, these tetradecamers remain catalytically inactive until binding of substrate peptide at the catalytic site (**Fig 5**). To date, all crystal structures of small-molecule-activator-bound hClpP have been resolved in a compact state and are largely interpreted as transition states due to a limited understanding of the hClpXP activation mechanism^21,24,44^. In light of our findings, it seems unlikely that the compact state is a transition state, but rather the resting state of the hClpP tetradecamer.

Our study highlights key regulatory adaptations that hClpXP has evolved to maintain mitochondrial proteostasis, which likely enhance the capacity of hClpXP to rapidly and efficiently respond to the dynamic mitochondrial environment. Consequently, our findings pinpoint regulatory facets that should be carefully considered in future investigations that seek to characterize the diverse roles of hClpXP within the mitochondrial space. We anticipate that our insights into assembly and activation of the hClpXP complex will lay the groundwork for impactful new studies exploring how these mechanisms are used to maintain mitochondrial proteostasis and cellular health.

## Materials and Methods

### hClpXP construct optimization

In our hands, the full-length hClpX protein (excluding the mitochondrial targeting sequence) did not form stable hexamers or form complexes with purified wild type hClpP. In pursuit of a hClpXP complex, we generated a hClpX construct lacking the flexible N-terminal Zinc binding domain, which is primarily involved in substrate recognition and adaptor binding^16,41^ and is routinely truncated for structural studies in bacterial systems^14,46^. Additionally, we removed the hClpX C-terminal extension, which is predicted to be disordered by AlphaFold (**Supplementary Fig. 1c**) and introduced a Walker B motif mutation (E359Q), which has been shown in bacterial ClpX and other AAA+ translocases to slow ATP hydrolysis and promote hexamer stability^11–13^.

This optimized hClpX construct purified as hexamers, but it did not readily form stable complexes with wildtype hClpP (**Supplementary Fig. 3a**). We next optimized hClpP construct to increase its affinity for hClpX. It was previously shown that removing the C-terminal extension from hClpP enhances hClpX binding^17^, but in our hands this construct did not form stable hClpXP complexes for cryo-EM structure determination. Introducing a proteolytically inactivating mutation into hClpP (S153A) promoted a marginal increase in hClpXP complex formation but was insufficient for single particle cryo-EM analysis (**Supplementary Fig. 3b**). We ultimately found that disuccinimidyl suberate (DSS) crosslinking^47^ stabilized the hClpXP construct for sample preparation and high-resolution cryo-EM analyses (**Supplementary Fig. 3c**). For simplicity, we refer to this optimized, crosslinked complex as hClpXP throughout the paper, unless otherwise noted.

### Protein expression and purification

The genes encoding for hClpX (residues 66-633) and hClpP (residues 58-277) were cloned into a PET28a expression vector containing a His_6_-SUMO tag at the N-terminus. Various hClpX variants (hClpX^I146-dCTE^, hClpX^I146-dCTE-dE^, hClpX^I146-dCTE-E396Q^, and hClpX^I146-dCTE-E396A^) and hClpP variants (hClpP^dCTE^ and hClpP^dCTE-S153A^) were generated using the Q5 Site-Directed Mutagenesis Kit from New-England Biolabs. The hClpX and hClpP constructs were transformed into BL21 Rosetta competent cells and grown overnight in LB medium supplemented with 50 µg/ml kanamycin and 25 µg/ml chloramphenicol. The overnight culture was then diluted 1:100 in fresh LB medium and incubated at 37 °C with a shaking speed of 225 rpm. When OD_600_ reached 0.6, protein expression was induced by adding isopropyl beta-D-thiogalactopyranoside (IPTG) at a final concentration of 0.5 mM, and the culture was shaken overnight at 16 °C. The following day, cells were harvested by centrifugation at 4500 rpm for 30 minutes, and the pellet was resuspended in lysis buffer (25 mM HEPES pH 7.5, 100 mM KCl, 400 mM NaCl, 10% glycerol, 0.1 M PMSF protease inhibitor). The cell suspension was sonicated, and the lysate was clarified by centrifugation at 35,000 rpm for 30 minutes. The supernatant containing the proteins was purified using Ni-NTA beads. The eluted proteins were cleaved with Ulp1 protease and dialyzed overnight in a buffer containing 25mM HEPES pH 7.5, 300 mM KCl, and 10% glycerol. To remove the His_6_-SUMO tag and Ulp1 protease, the cleaved proteins were passed through Ni-NTA beads again. The proteins were subsequently further purified using size exclusion chromatography (SEC) on a Superdex 200 10/300 GL column. For hClpX, the SEC buffer contains 25 mM HEPES pH 7.5, 300 mM KCl, 10% glycerol, 2 mM magnesium, and 1 mM DTT. hClpP was purified using SEC buffer with containing 25 mM HEPES pH7.5, 150 mM KCl, 5% glycerol, and 1 mM DTT. Purified proteins were concentrated, aliquoted, flash frozen in liquid nitrogen, and stored at −80 °C.

### Sample preparation for electron microscopy

For all three cryo-EM structures presented in this paper, 4 µl of sample was applied onto 300 mesh R1.2/1.3 UltraAuFoil Holey Gold grids (Quantifoil), which were plasma cleaned for 15 seconds using a Pelco glow discharge cleaning system (TED PELLA, INC.) with atmospheric gases at 15 mA. The grids were manually blotted using Whatman #1 blotting paper for 6 seconds before being plunged frozen in liquid ethane in a 4 °C cold room with 90% humidity (all subsequently described vitrification of samples were performed using the same methodology).

To prepare hClpXP sample, hClpX^I146-dCTE-E396Q^ (50 µM) and hClpP^dCTE-S153A^ (8µM) were in incubated in a buffer containing 25 mM HEPES (pH 7.5), 150 mM KCl, 5 mM ATP, and 5 mM MgCl_2_ for 5 minutes at room temperature. The sample was then crosslinked with 160 µM disuccinimidyl suberate (DSS) (Thermo Fisher A39267) for 30 minutes at room temperature, and the reaction was quenched with 40 µM Tris (pH 7.5). The crosslinked sample was concentrated and injected onto a Superose 6 increase 10/300 GL column in SEC buffer (25 mM HEPES, 150 mM KCl, 5% glycerol, 2 mM MgCl2, 2 mM ATP). Fractions corresponding to the crosslinked hClpXP complexes were collected and concentrated 15-fold. The concentrated sample was then applied to the cryoEM grid and manually blotted before being plunged frozen in liquid ethane.

For the apo hClpP cryo-EM structure, 4 µl of 15 mg/ml of hClpP (WT) in a buffer containing 25 mM HEPES, 150 mM KCl, 2 mM DTT, and 4mM fluorinated fos-choline detergent was applied onto the cryo-EM grid and immediately blotted before being plunged into liquid ethane. For the bortezomib-bound hClpP structure, a 10X molar excess of bortezomib (Sigma-Aldrich) was added to 15 mg/ml of hClpP and incubated on ice for 30 minutes. Just before applying the sample to the cryo-EM grid, n-Decyl-beta-Maltoside (DM) detergent at final 0.08% concentration was added to the sample to help mitigate preferred orientation of hClpP in vitrified ice.

### Cryo-EM data collection and processing

For all datasets, cryo-EM data were collected on a Thermo-Fisher Talos Arctica transmission electron microscope operating at 200 kV with parallel illumination conditions^48,49^. Micrographs were acquired with a Gatan K2 Summit direct electron detector with a total electron exposure of 50 e^−^/Å^2^. All subsequent image processing was performed using cryoSPARC^50^. Detailed processing workflows and dataset-specific parameters are provided in **Supplementary Figs. 4, 10, 13** and **Table 1**.

For the hClpXP complex cryo-EM dataset, Leginon data collection software^51^ was used to collect 10,333 micrographs at 45K nominal magnification (0.91 Å /pixel^−1^), with a nominal defocus range from −0.23 µM to −3.3 µm. Stage movement was used to target the center of either 4 or 16 1.2 µm holes for focusing, and image shift was utilized to acquire high magnification images. The Appion image processing wrapper^52^ was used to run MotionCor2^53^ for micrograph frame alignment and dose-weighting in real-time during data collection. Subsequent image processing was done in cryoSPARC. The contrast transfer function (CTF) parameters were estimated using patch CTF estimation (multi). Template-based particle picking, using representative 2D class averages generated from 1,175 particles selected with blob picking using a diameter of 200 Å to 320 Å, resulted in approximately 2.07 million particles. Particles were extracted from the motion-corrected and dose-weighted micrographs with a box size of 352 pixels. These particles were classified into 100 2D classes using default parameters, and 2D classes (containing 1,940,578 particles) displaying high-resolution secondary features were selected to generate four reference-free 3D Abinitio models. The initial volumes generated from Ab-initio were used for heterogenous refinement to classify the particles into four classes with C1 symmetry. The class containing well-resolved structural features, corresponding to 1,321,170 particles, was chosen for non-uniform refinement of hClpP with C1 symmetry, resulting in a final reconstruction with a reported GSFCS at 2.91 Å. For hClpX, the same particle set underwent additional 3D classification (six classes) with a mask corresponding to hClpX (O-EM learning rate init: 0.2, initialization mode: PCA; PCA/simple: particles per reconstruction: 750; PCA: number of reconstructions: 75; Force hard classification; class similarity: 0.1). Two good classes were combined and subjected to a final round of non-uniform refinement with C1 symmetry. Subsequently, the 3D volume was re-aligned centering on hClpX, and the particles re-extracted with the centered coordinates were local refined with a mask around hClpX, resulting in a final resolution of 3.11 Å based on the FSC at a cutoff of 0.143.

For the apo hClpP (WT) structure, a total of 14,989 micrographs were collected at 150K nominal magnification (0.94 Å /pixel^−1^) with a nominal defocus range from −0.3 µm to −3.5 µm using the EPU software. A maximum image shift of 2 µm was used to acquire high magnification images. Dose-weighting, motion correction, and CTF estimation were performed within cryoSPARC live. Template-based particle picking, using representative 2D class averages generated from 4,710 particles selected with blob picking (diameter of 70 to 100 Å), resulted in ~2.7 million particles. Two rounds of 2D classification into 100 class averages using default parameters were performed to remove junk particles, with 935,410 particles selected for Ab-initio and iterative hetero-refinement (four classes). The class with the best-resolved structural features (corresponding to 501,826 particles) was used for non-uniform refinement (C1 symmetry), yielding an hClpP reconstruction at 3.63 Å resolution. Subsequent global and local CTF refinement, and another round of non-uniform refinement slightly improved the resolution to 3.61 Å. 3D classification into 10 classes were performed, with particles from eight good classes used for the final round of non-uniform refinement (C1 symmetry), resulting a final reported GSFCS resolution of 3.52 Å.

For the bortezomib-bound-hClpP dataset, 3,972 micrographs were collected using EPU at 150K nominal magnification (0.94 Å /pixel^−1^) with a nominal defocus range from −0.1 µm to −3.28 µm. Image shift was used to acquire high magnification images as for the prior dataset. Dose-weighting, motion correction, and CTF estimation were performed within cryoSPARC live. Blob picking of the CTF estimated images using a particle diameter of 100 Å to 150 Å resulted in ~1.9 million particles. Two rounds of 2D classification were performed to remove junk particles, leaving 545,529 particles for Ab-initio and subsequent hetero-refinements (4 classes). The class with the best-resolved structural features (corresponding to 224,782 particles) was used for non-uniform refinement (C1 symmetry), producing hClpP reconstruction at 2.67 Å resolution. Further global and local CTF refinement slightly increased the resolution to 2.63 Å with C1 symmetry and 2.28 Å with D7 symmetry.

### Atomic model building and refinement

For the hClpXP complex, the AlphaFold model of hClpX and the hClpP crystal structure (PDB 6BBA) were used as starting models. For the bortezomib-bound hClpP, the hClpP crystal structure (PDB 1TG6) was used as a starting model. For the hClpP apo structure, the hClpX-bound hClpP model was used. The models were docked into the 3D reconstruction using UCSF ChimeraX^54^. Manual model building and realspace structural refinement were performed with Coot^55^ and Phenix^56^, respectively. Phenix real space refinement includes global minimization, rigid body, local grid search, atomic displacement parameters, and morphing for the first cycle. It was run for 100 iterations, five macro cycles, with a target bonds RMSD of 0.01 and a target angles RMSD of 1.0. The refinement settings also include the secondary structure restraints, Ramachandran restraints. ChimeraX plug-in ISOLDE^57^ was used to manually fix any Ramachandran outliers, rotamers, and clashes not fixed by Phenix. For the bortezomib-bound hClpP, a covalent linkage between bortezomib and hClpP S153 was made using Phenix. Figures for publication were prepared using UCSF ChimeraX. Table 1 includes data collection, refinement, and validation statistics.

### ATPase activity assay

The ATPase activity of different hClpX variants were carried out in a black flat-bottom 96-well plate in a final volume of 100 µl. 0.3 µM of hClpX_6_ proteins was pre-incubated at 37 °C for 5 min in the activity buffer (50 mM HEPES pH 7.5, 75 mM KCl, 5 mM MgCl2, and 1mM DTT, 1.6 mM NADH (Caymen Chemical; 606-69-8), 4 mM phosphoenolpyruvate (1PlusChem; 5541-93-5), 120 U/ml lactate dehydrogenase (Worthington Chemical; LS002755), 40 U/ml pyruvate kinase (Sigma; 10128155001).) The reaction was initiated by adding 5 mM ATP. Absorption at 340 nm was measured at 37 °C using a TECAN plate reader.

### Degradation assays

The ClpXP-mediated degradation of FITC-casein was carried out in a black flat-bottom 96-well plate in a final volume of 100 µl. 1 µM ClpX and 0.5 µM ClpP were pre-incubated at 37 °C for 5 min in activity buffer containing 50 mM HEPES pH7.5, 75 mM KCl, 5 mM MgCl2, 5 mM ATP, and 1 mM DTT. 1 µM FITC-casein (Sigma Aldrich; C3777), which had also been pre-incubated at 37 °C, was added to initiate the degradation reaction. An increase of fluorescence resulting from FITC molecules was measured (excitation 485 nm, emission 535 nm) using a TECAN plate reader.

## Acknowledgements

We thank Jeff Mindrebo and the members of the Lander lab for helpful discussion. We thank Will Lessin at the Scripps Research Electron Microscopy Facility for microscopy support. We thank Charles Bowman and J.C. Ducom at Scripps Research High Performance Computing for computational support. This work is supported by a grant from the National Institutes of Health (NIH) NS095892 to G.C.L. and a National Science Foundation predoctoral fellowship to W.C. Cryo-EM data collection used equipment supported by NIH grant S10OD032467.

## Data availability

Cryo-EM maps and associated atomic models were deposited to the Electron Microscopy Databank (EMDB) and the Protein Databank (PDB), respectively, with the following EMDB and PDB IDs: hClpXP (focused alignment of ClpX): EMD-47226, 9DVY; hClpXP (focused alignment of ClpP): EMD-47232, 9DW0; hClpP heptamer: EMD-47233, 9DW1; hClpP tetradecamer bound to bortezomib: EMD-47234, 9DW3.

## Author Contributions

J.Y. and G.C.L. initially conceptualized the project. W.C. and G.C.L. led the project; W.C. generated the constructs, purified the proteins, and performed all cryo-EM structure determination, model building and refinement, and mechanistic interpretation with guidance from J.Y. and G.C.L.; W.C. and G.C.L. wrote the manuscript.

## Supplementary Figures

**Supplementary Figure 1.**
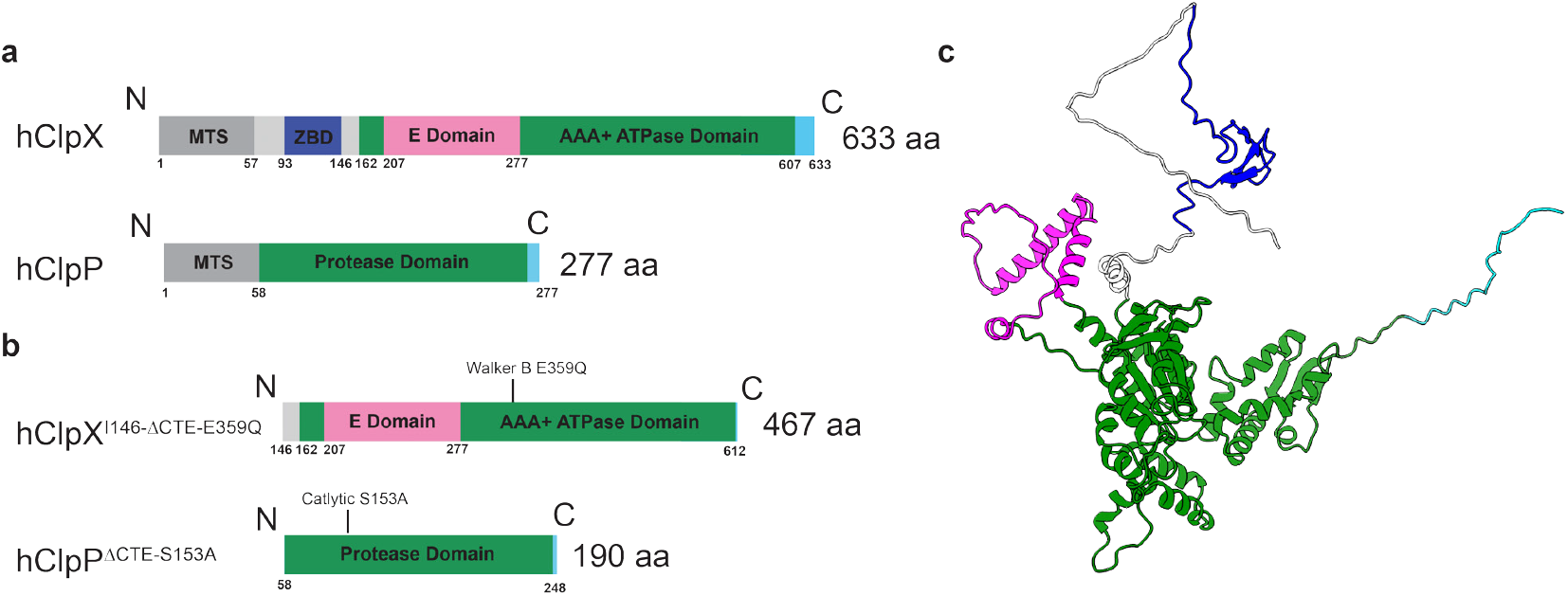
hClpXP construct optimization for cryo-EM. (**a**) Schematic of domain organization of wild type hClpX and hClpP. (**b**) Domain organization of hClpX and hClpP constructs used for this cryo-EM study. (**c**) AlphaFold structure of monomeric full-length hClpX (lacking the MTS), with different domains colored corresponding to the domain architecture.

**Supplementary Figure 2.**
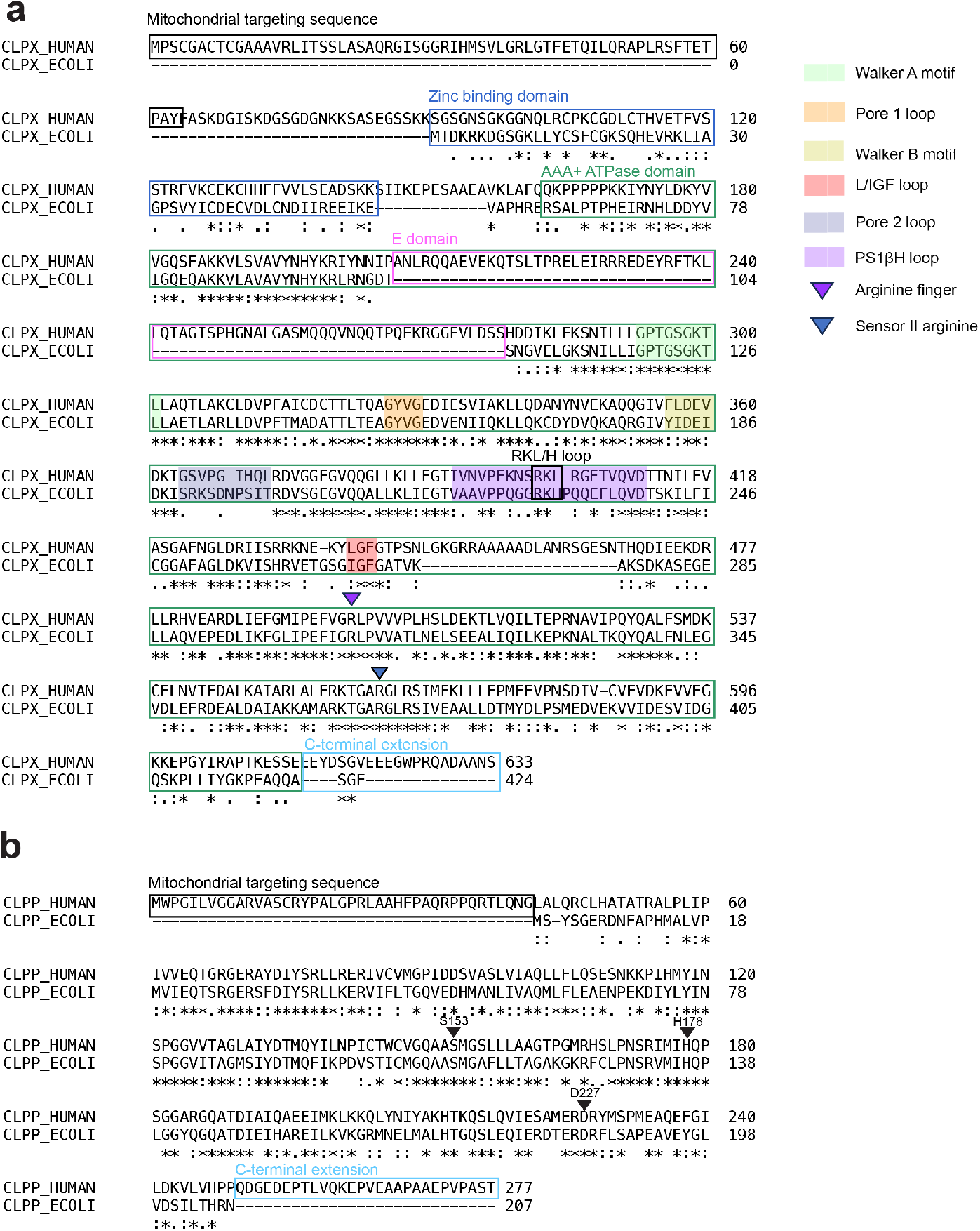
Sequence alignment of human and *E*.*coli* ClpXP. (**a**) ClustalW alignment of the human and *E*.*Coli* ClpX sequences. Asterisks denotes strict conservation, and one or two dots indicate high sequence similarity. Key domains are denoted with boxes and the conserved motifs are highlighted with solid colors. The arginine finger and the sensor II arginine are denoted with arrows. hClpX features a unique insertion known as the E-loop (pink box), as well as additional residues at the C-terminus (cyan box). (**b**) ClustalW alignment of the human and *E*.*Coli* ClpP sequences. The catalytic residues are denoted with arrows. Like hClpX, hClpP contains additional residues at the C-terminus, referred to as the C-terminal extension.

**Supplementary Figure 3.**
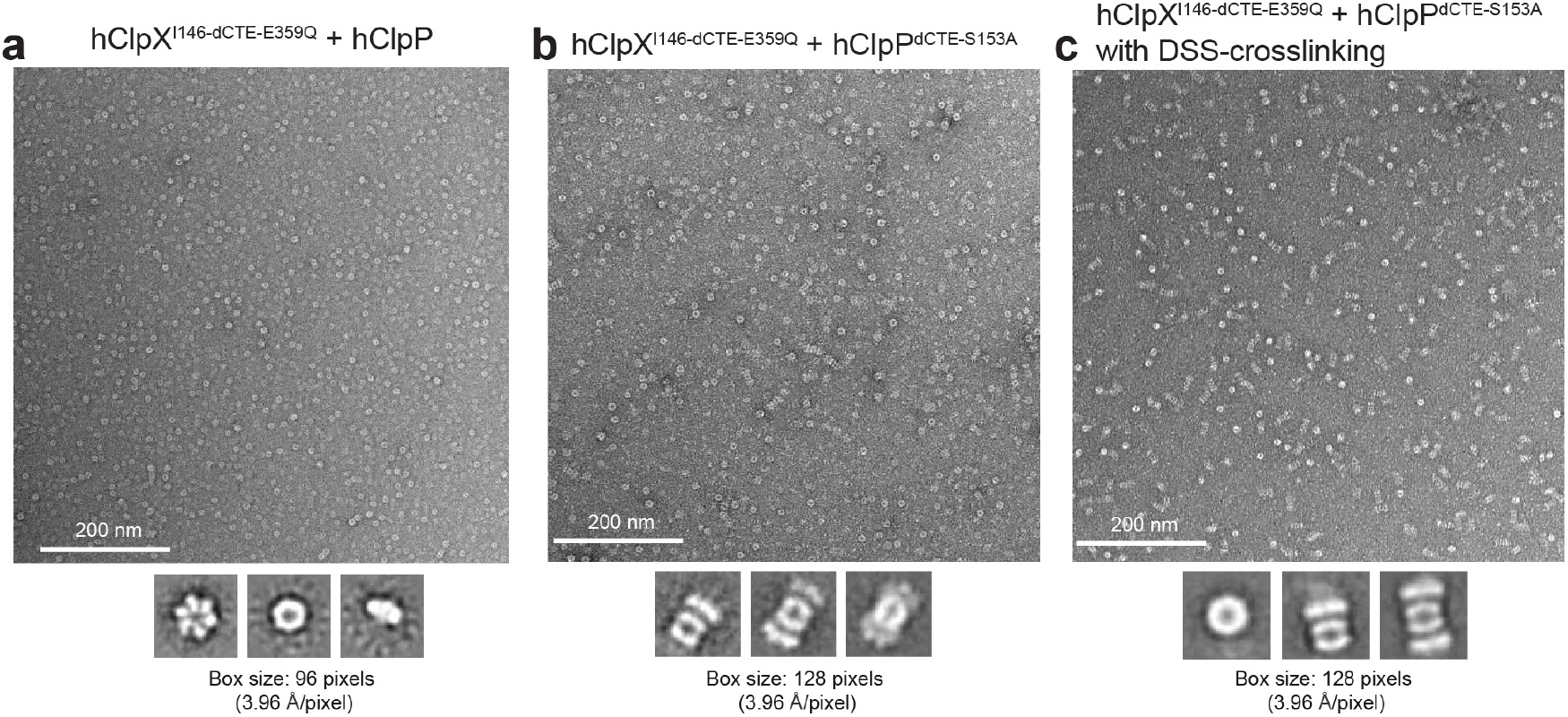
Negative staining analyses of hClpXP complexes. (**a**) Representative negative stain micrograph of a sample mixture containing hClpX^I146-dCTE-E359Q^, hClpP, ATP, and MgCl_2_. The interaction between hClpX^I146-dCTE-E359Q^ and hClpP are not stable, as no hClpXP complexes are observed. Below the micrograph, 2D averages of the hexameric ClpX, and heptameric ClpP are shown. (**b**) Representative micrograph of a hClpX^I146-dCTE-E359Q^ and hClpP^dCTE-S153A^ in the presence of ATP and MgCl_2_. Removal of the C-terminal extension and the introduction of the S153A mutation increases hClpP’s affinity for hClpX. Some hClpXP complexes, including tetradecameric hClpP with either single or double-capped hClpX, are observed, as shown in the 2D average below. However, the hClpXP complex represented a minority of the observed particles. (**c**) Representative micrograph and 2D averages of DSS-crosslinked hClpX^I146-dCTE-E359Q^ and hClpP^dCTE-S153A^, showing that the hClpXP complex makes up the majority of the particles.

**Supplementary Figure 4.**
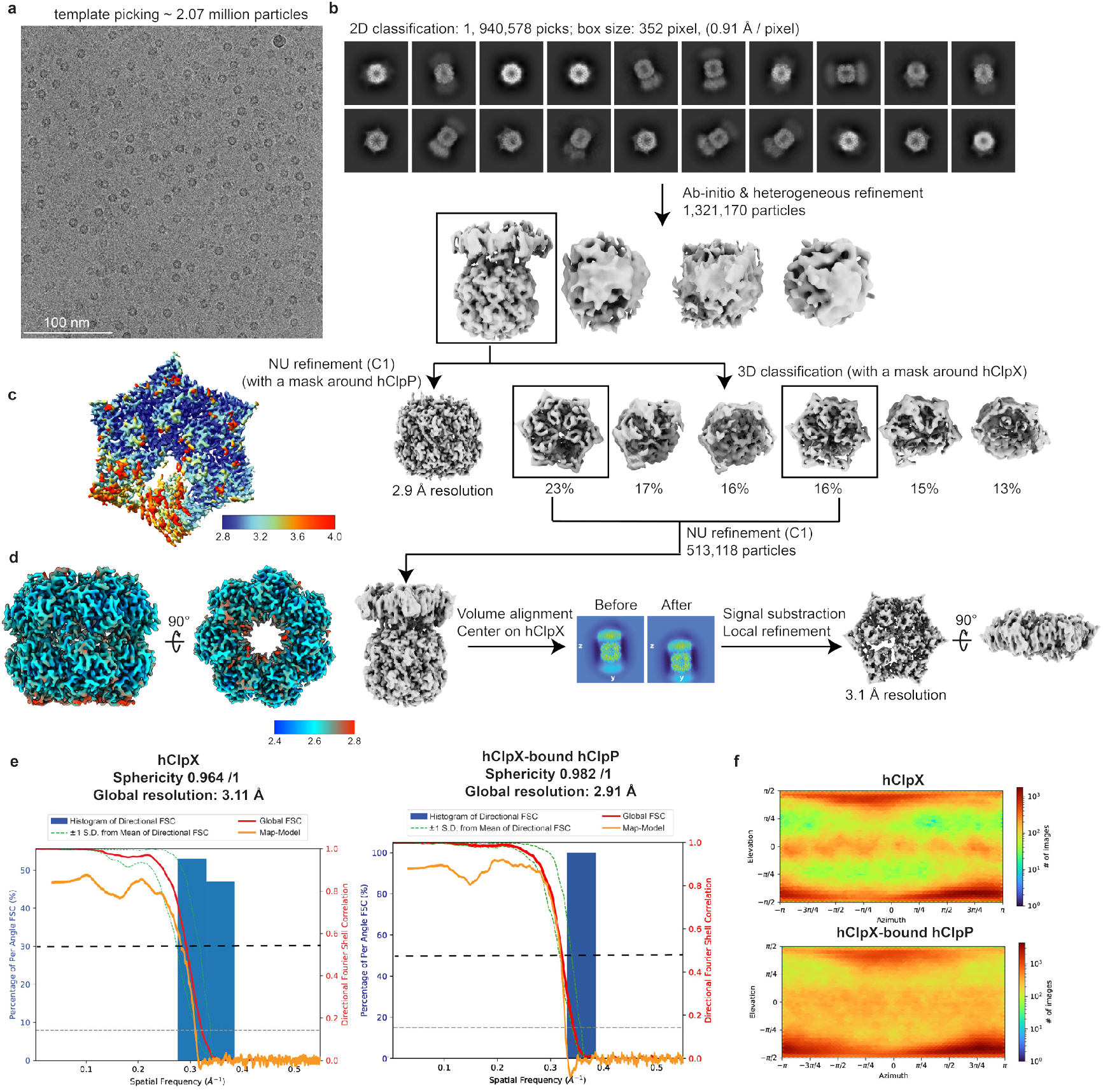
hClpXP cryo-EM processing workflow. (**a**) Representative motion-corrected, dose-weighted cryo-EM micrograph of vitrified hClpXP. (**b**) CryoSPARC processing scheme used to generate 3D reconstructions of hClpP and hClpX. (**c**) Local resolution map of hClpX calculated from cryoSPARC. (**d**) Local resolution map of hClpX-bound hClpP calculated from cryoSPARC. (e) Global FSC and map-to-model FSC overlaid on the directional resolution histogram from the 3DFSC server^45^. (**f**) Viewing distribution plot corresponding to focused hClpX and hClpP reconstructions.

**Supplementary Figure 5.**
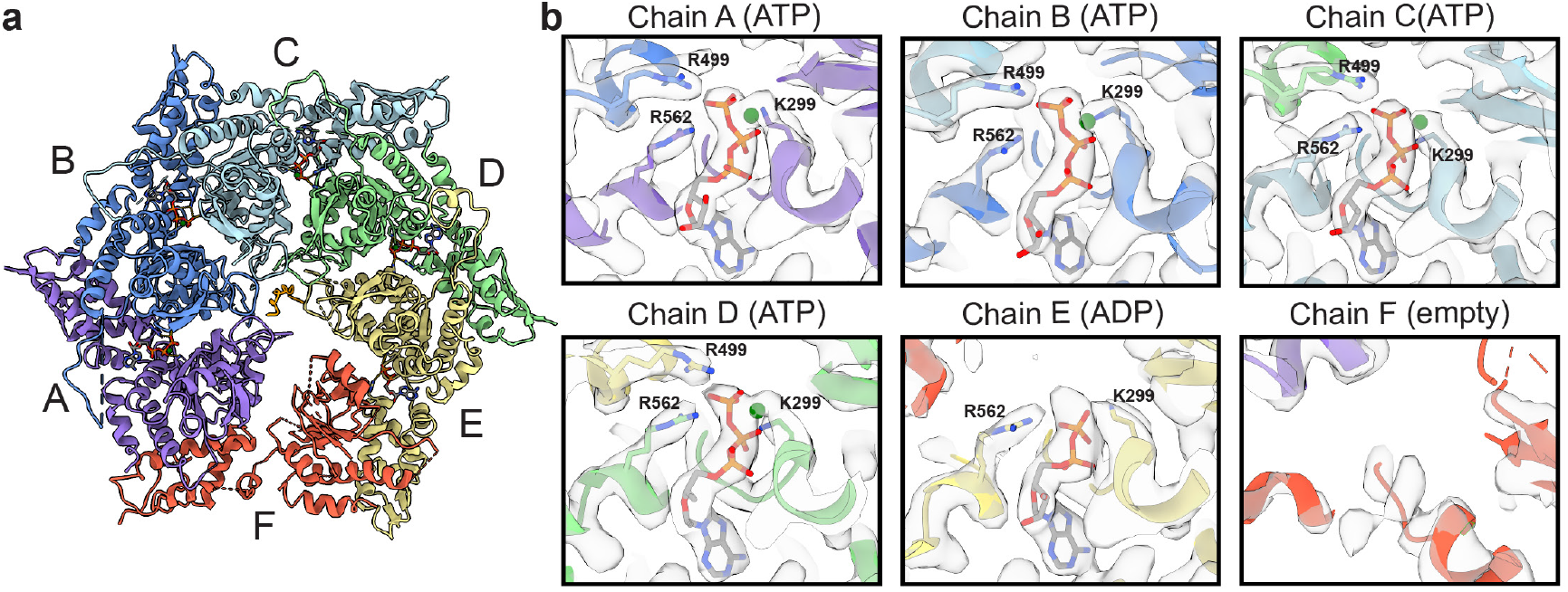
hClpX nucleotide density. (**a**) Ribbon representation of the atomic model of hClpX. (**b**) Nucleotide-binding sites for chains A-F, with key interacting residues shown along with experimental density maps corresponding to the bound nucleotides shown as transparent surfaces.

**Supplementary Figure 6.**
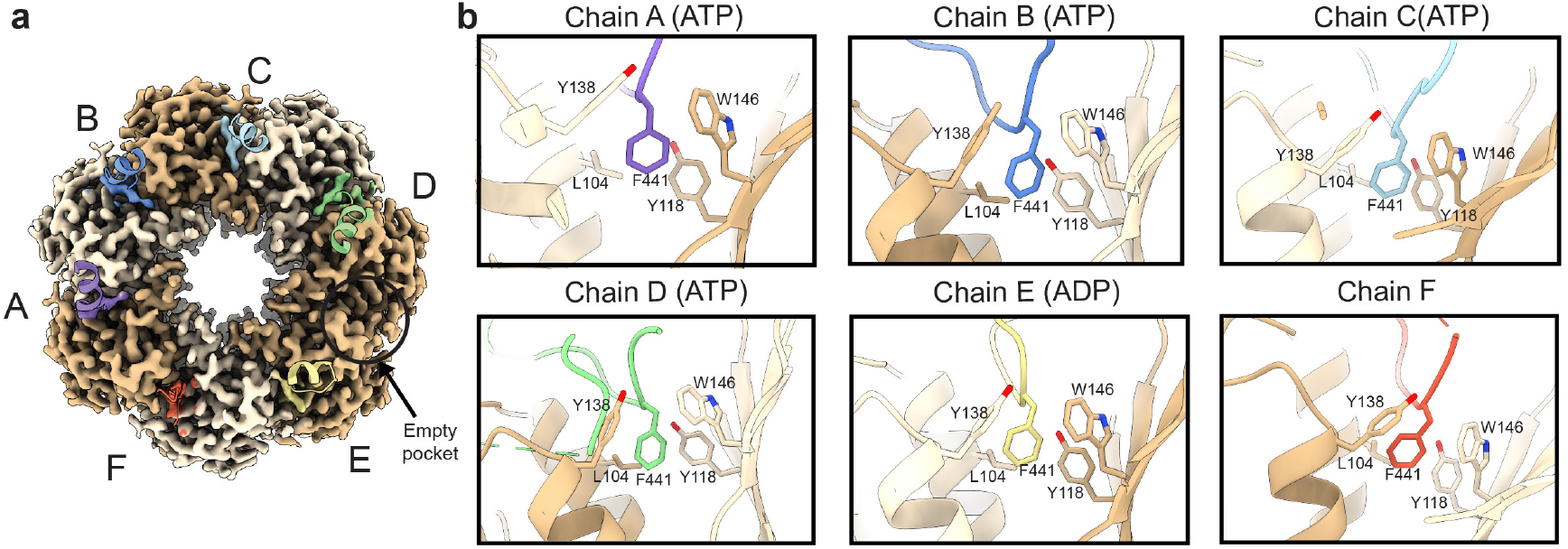
hClpX LGF loop and hClpP hydrophobic pocket interactions. (**a**) Axial view of hClpP density showing the hClpX LGF loops bound to the hydrophobic pockets on hClpP. The LGF loop of hClpX is depicted as a ribbon, while the EM density of the interacting residues shown and correspondingly colored. (**b**) Phe441 of the 6 LGF loops are shown interacting with the hydrophobic clefts of hClpP, with the interacting residues labeled.

**Supplementary Figure 7.**
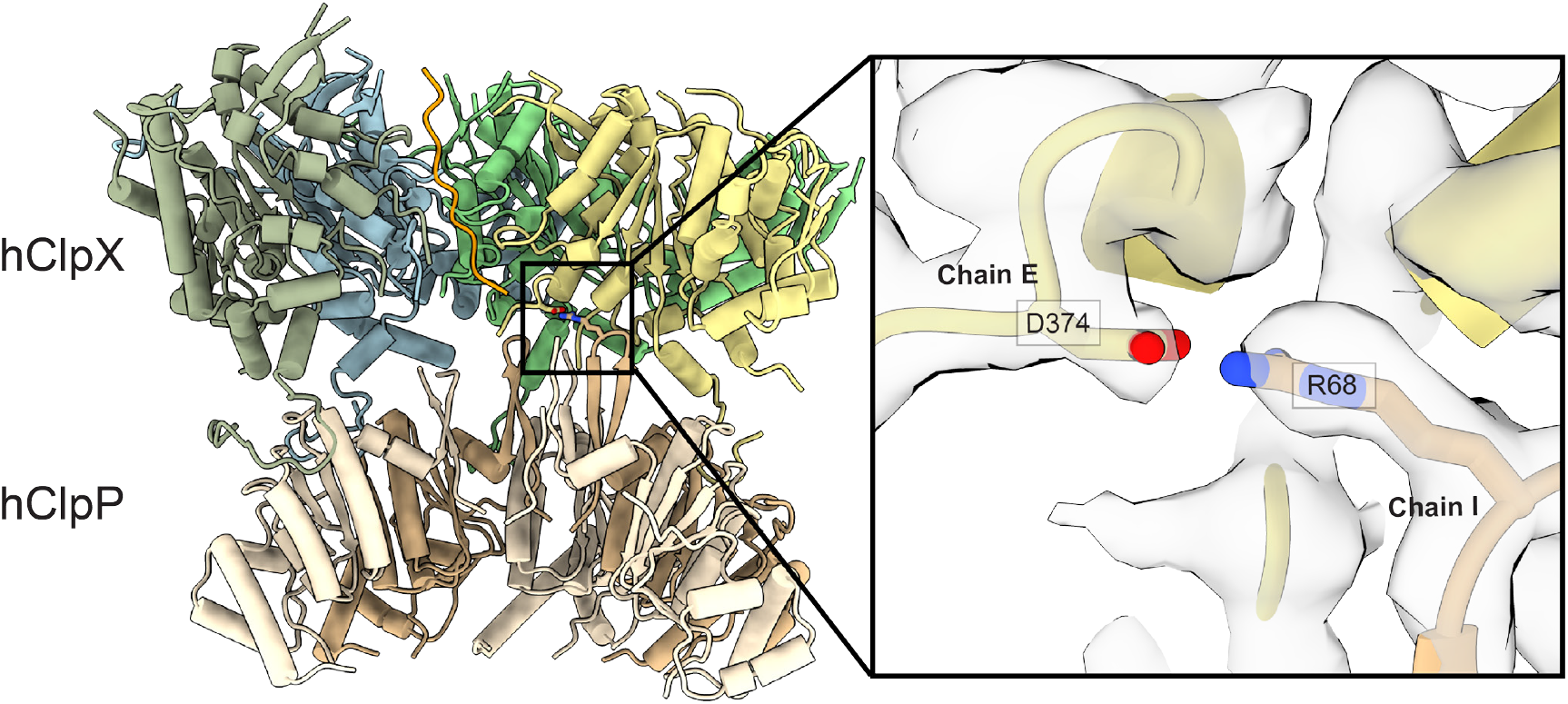
The interaction interface between hClpX and hClpP. hClpXP model highlighting the interface between hClpX and hClpP. The inset shows D374 from Chain E interacting with R68 at the top of N-terminal β-hairpin from chain I, with the experimental density shown as a transparent surface.

**Supplementary Figure 8.**
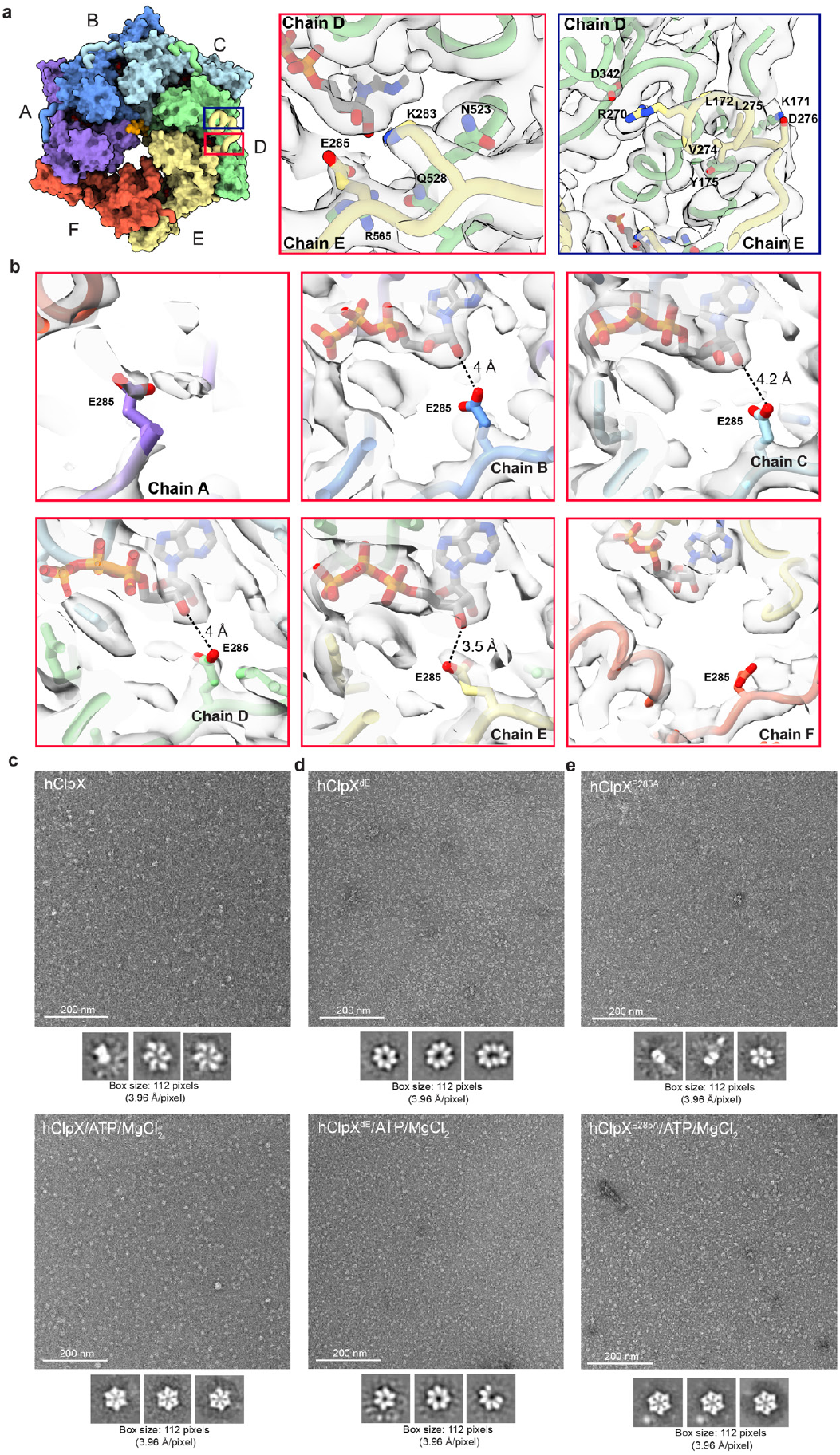
The E-loop facilitates hClpX hexamer formation. (**a**) A molecular surface representation of the hClpX hexamer with each subunit colored distinctly is shown on the left. The E-loops are not shown as a surface and are instead shown using a thick ribbon representation. The panels on the right show the density for the interacting residues within the E loop, which correspond to the regions denoted by the boxes shown on the hexamer. (**b**) The putative interaction between bound nucleotide and E285 from the E-loops in the different ClpX subunits (A-F) are shown as an atomic model with the experimental density shown as a transparent surface. (**c-e**) Representative negative staining micrographs and 2D averages of hClpX, hClpX^dE^, or hClpX^E285A^ in the absence or presence of the exogenous ATP and MgCl_2_. hClpX^dE^ forms higher order oligomers, and binding of ATP/MgCl_2_ can promote formation of loosely organized hClpX hexamers as well as other oligomers.

**Supplementary Figure 9.**
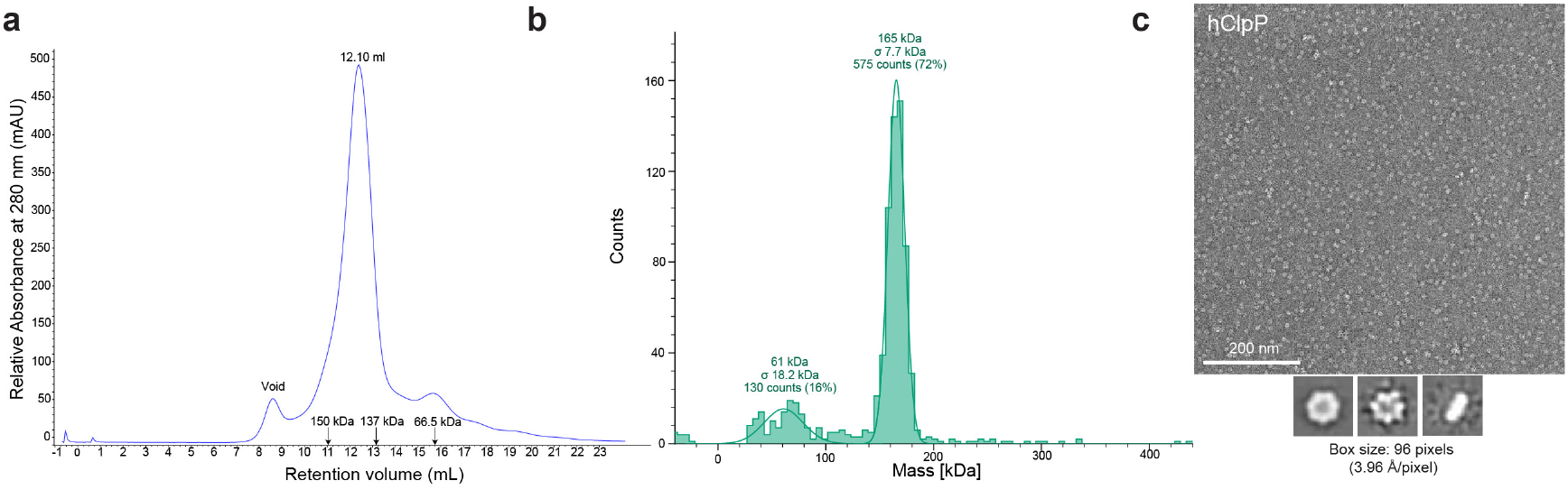
hClpP is a stable heptamer in solution. (**a**) A size exclusion chromatography trace of hClpP shows that the oligomer elutes at a fraction corresponding to a heptamer species, as determined by comparison with protein molecular weight standards. **b**, Mass photometry of hClpP shows that the oligomer is primarily heptameric. **c**, Representative negative staining image and 2D averages of hClpP heptamer.

**Supplementary Figure 10.**
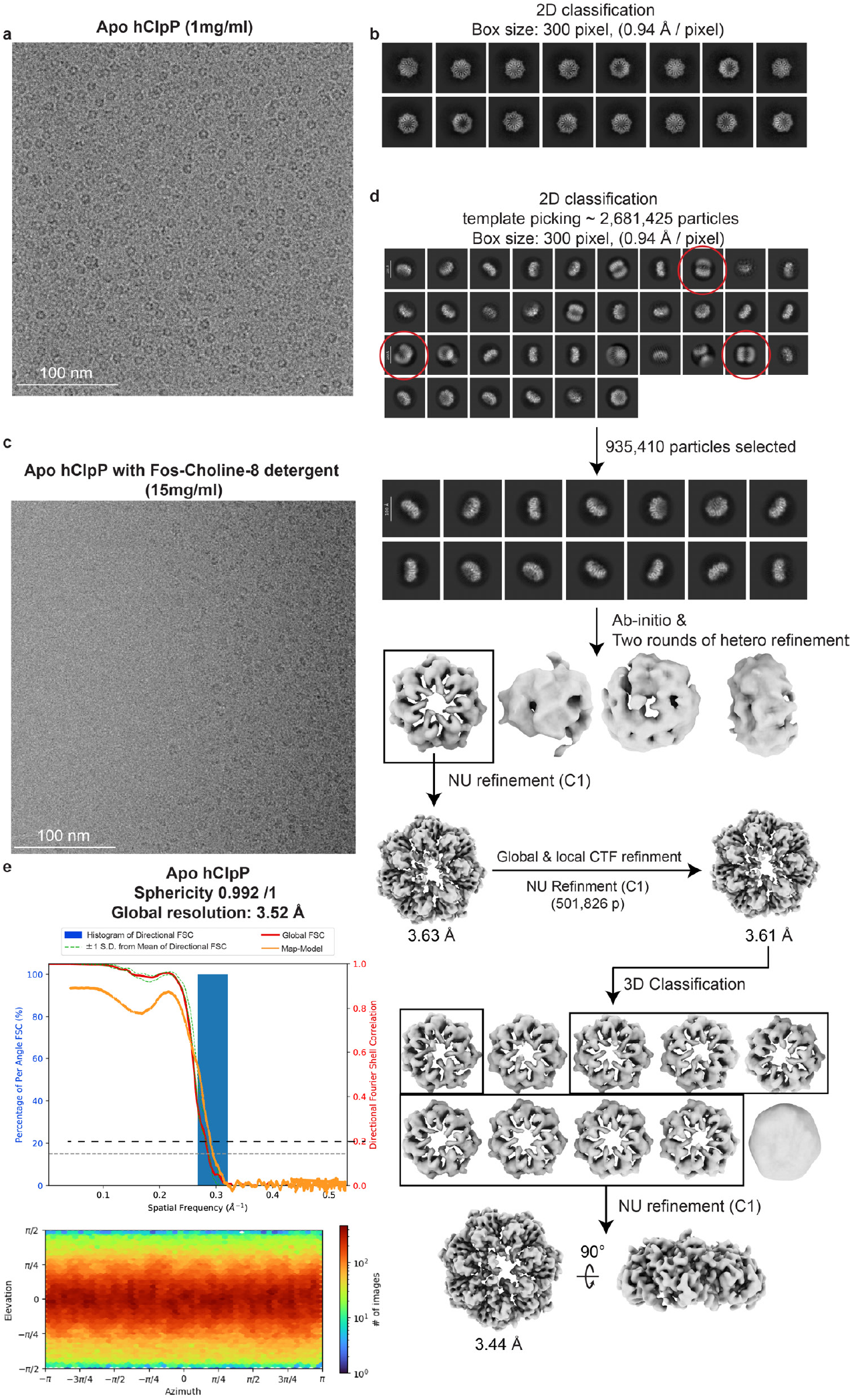
(previous page). Apo hClpP cryo-EM processing workflow. (**a**) Representative motion-corrected, dose-weighted cryo-EM micrograph of vitrified 1 mg/ml apo hClpP. (**b**) Representative 2D averages showing strong preferred axial orientation of hClpP. (**c**) Representative motion-corrected, dose-weighted cryo-EM micrograph of 15 mg/ml hClpP with fluorinated fos-choline-8 detergent. **(d)** CryoSPARC processing scheme used to generate 3D reconstruction of the apo hClpP. (**e**) Local resolution map of apo hClpP calculated from cryoSPARC. (**f**) Global FSC and map-to-model FSC overlaid on the directional resolution histogram from the 3DFSC server^45^. (**g**) Viewing distribution plot of the apo hClpP reconstruction.

**Supplementary Figure 11.**
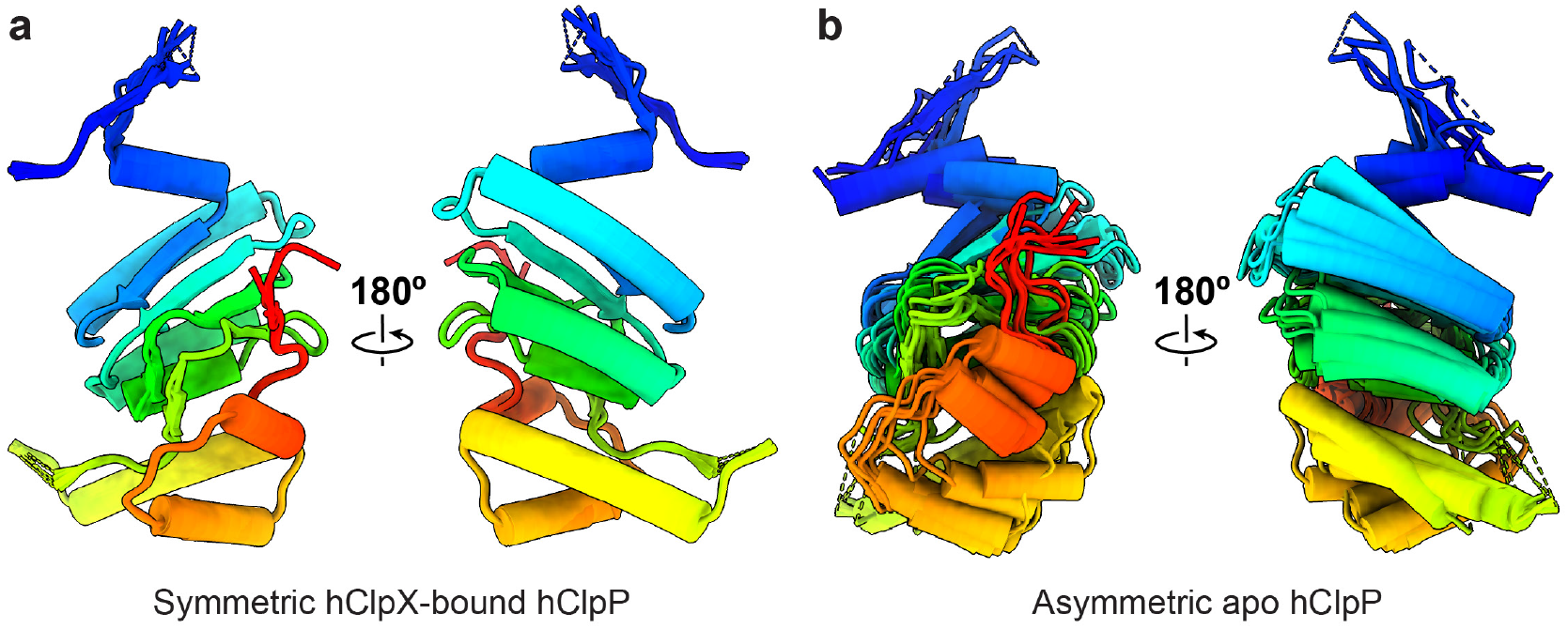
Overlay of hClpP subunits in tetradecameric and heptameric oligomers. (**a**) Seven-fold symmetry parameters were used to overlay individual subunits of ClpP from one ClpP ring of the ClpXP complex refined without imposed symmetry. The subunits overlay perfectly, demonstrating the symmetric arrangement of the subunits in the context of the tetradecamer. (**b**) The same operation was applied to the subunits of the asymmetric apo hClpP heptamer, showing the distinct positions of the individual subunits.

**Supplementary Figure 12.**
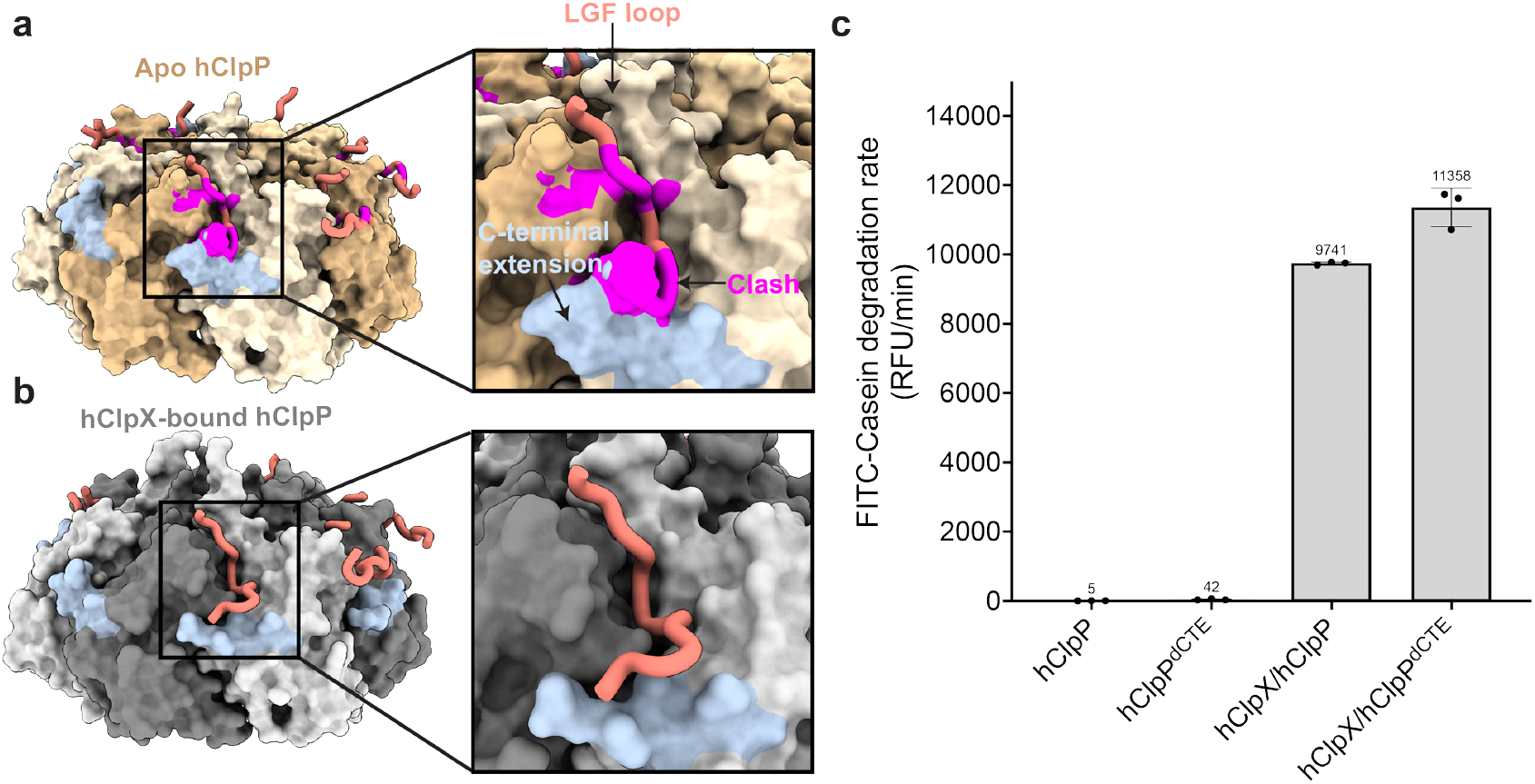
The C-terminal extension of apo hClpP sterically hinders hClpX LGF binding. **(a)** Surface representation of apo hClpP (top) and (**b**) hClpX-bound hClpP (bottom), with the C-terminal extension highlighted in light steel blue and the LGF loops shown using a thick ribbon representation, colored salmon. In the apo hClpP structure, the C-terminal extension sterically clashes with the LGF loops from hClpX. To accommodate hClpX binding, the C-terminal extension shifts downward, as observed in the hClpX-bound hClpP structure. (**c**) FITC-casein degradation assays show that neither hClpP or hClp^dCTE^ alone is capable of degrading casein substrates. Degradation activity is observed only in the presence of hClpX. Removing the C-terminal extension from hClpP marginally enhances its enzymatic activity.

**Supplementary Figure 13.**
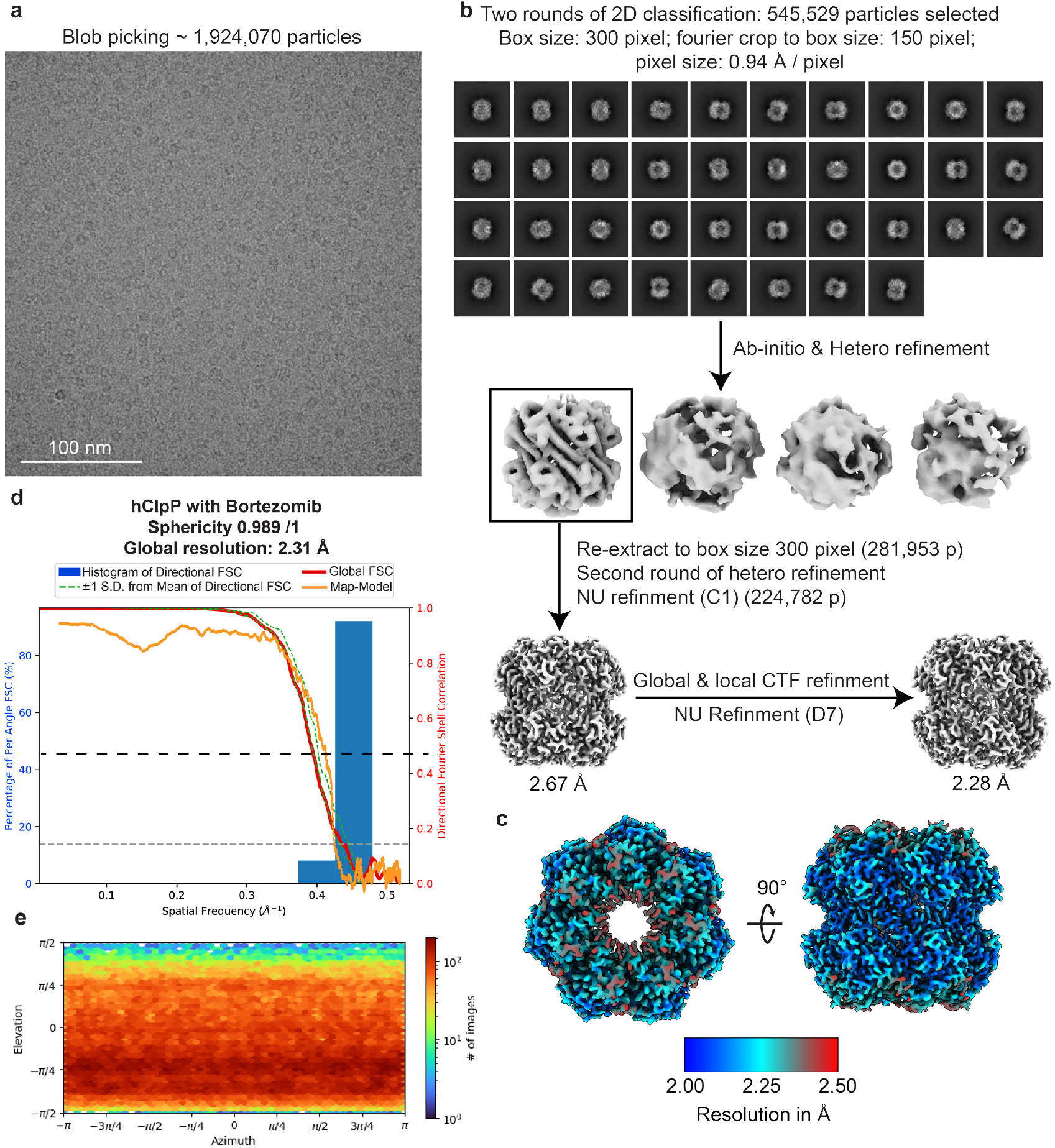
Bortezomib-bound hClpP cryo-EM processing workflow. (**a**) Representative motion-corrected, dose-weighted cryo-EM micrograph of vitrified hClpP bound to bortezomib. (**b**) CryoSPARC processing scheme used to generate 3D reconstruction of the bortezomib-bound hClpP. (**c**) Local resolution map of apo hClpP calculated from cryoSPARC. (**d**) Global FSC and map-to-model FSC overlaid on the directional resolution histogram from the 3DFSC server^45^. (**e**) Viewing distribution plot corresponding to bortezomib-bound hClpP.

**Supplementary Figure 14.**
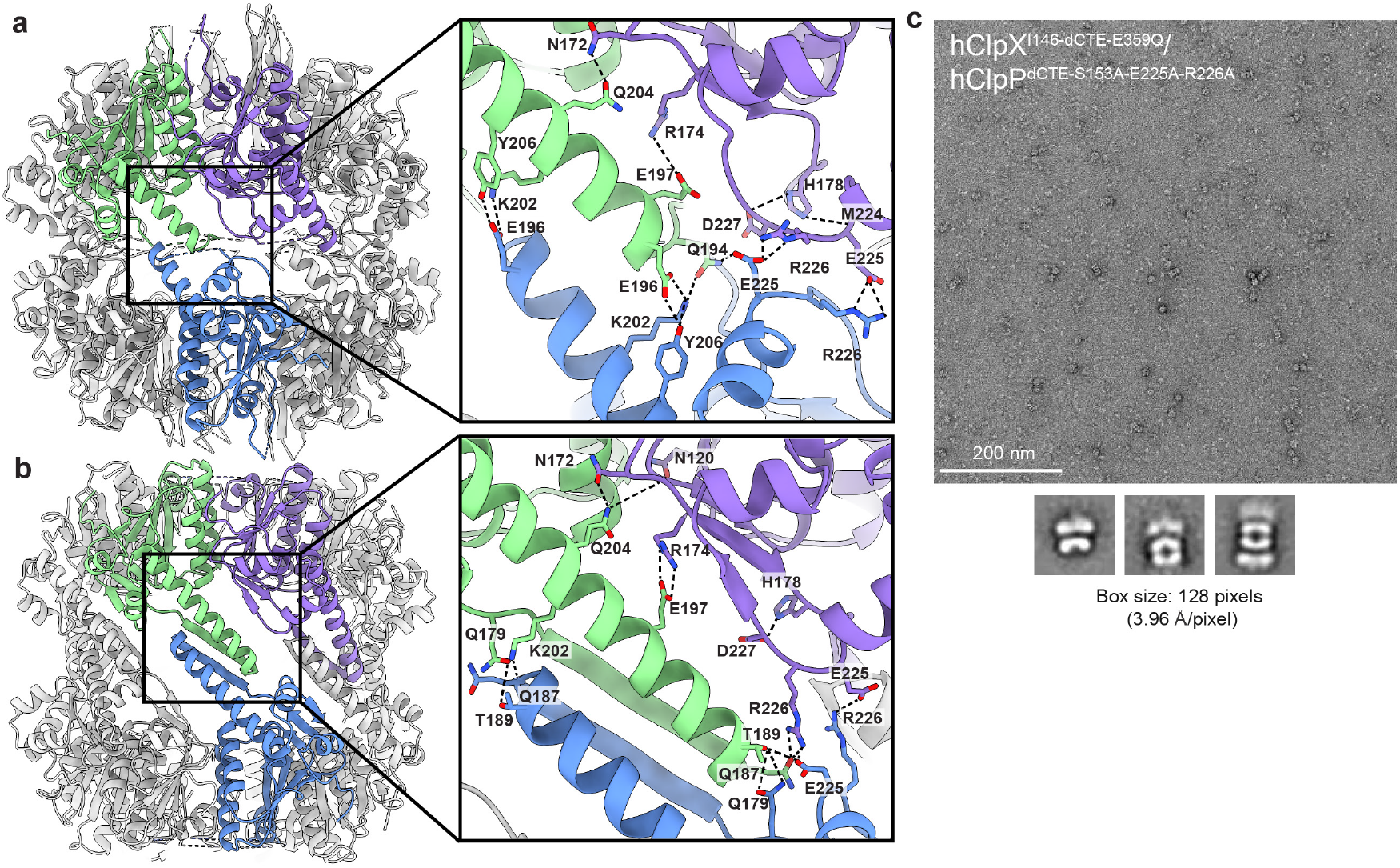
Key interactions at the hClpP-hClpP interface of compact and extended hClpP tetradecamer. (**a**) Atomic models of the hClpX-bound compact hClpP and (**b**) bortezomib-bound extended hClpP reveal a network of interacting residues at the hClpP-hClpP interface. (**c**) Representative micrograph and 2D averages of crosslinked-hClpX^I146-dCTE-E359Q^ and hClpP^dCTE-S153A-E225A-R226A^. Mutations of oligomeric sensors E225 and R226 destabilize but do not abolish ClpP tetradecamer formation.

## References

1. Osellame, L.D., Blacker, T.S. & Duchen, M.R. Cellular and molecular mechanisms of mitochondrial function. Best Pract Res Clin Endocrinol Metab 26, 711–23 (2012).

2. Kuznetsov, A.V. & Margreiter, R. Heterogeneity of mitochondria and mitochondrial function within cells as another level of mitochondrial complexity. Int J Mol Sci 10, 1911–1929 (2009).

3. Tait, S.W.G.G. D. R.,. Mitochondrial regulation of cell death. Cold Spring Harb Perspect Biol 5(2013).

4. Jadiya, P.T. D.,. Mitochondrial protein quality control mechanisms. Genes (Basel) 11(2020).

5. Javadov, S., Kozlov, A.V. & Camara, A.K.S. Mitochondria in Health and Diseases. Cells 9(2020).

6. Kardon, J.R., Moroco, J.A., Engen, J.R. & Baker, T.A. Mitochondrial ClpX activates an essential biosynthetic enzyme through partial unfolding. Elife 9(2020).

7. Kardon, J.R. et al. Mitochondrial ClpX Activates a Key Enzyme for Heme Biosynthesis and Erythropoiesis. Cell 161, 858–67 (2015).

8. Kasashima, K., Sumitani, M. & Endo, H. Maintenance of mitochondrial genome distribution by mitochondrial AAA+ protein ClpX. Exp Cell Res 318, 2335–43 (2012).

9. Seo, J.H. et al. The Mitochondrial Unfoldase-Peptidase Complex ClpXP Controls Bioenergetics Stress and Metastasis. PLoS Biol 14, e1002507 (2016).

10. Mabanglo, M.F. & Houry, W.A. Recent structural insights into the mechanism of ClpP protease regulation by AAA+ chaperones and small molecules. J Biol Chem 298, 101781 (2022).

11. Ripstein, Z.A., Vahidi, S., Houry, W.A., Rubinstein, J.L. & Kay, L.E. A processive rotary mechanism couples substrate unfolding and proteolysis in the ClpXP degradation machinery. Elife 9(2020).

12. Fei, X. et al. Structures of the ATP-fueled ClpXP proteolytic machine bound to protein substrate. Elife 9(2020).

13. Gatsogiannis, C., Balogh, D., Merino, F., Sieber, S.A. & Raunser, S. Cryo-EM structure of the ClpXP protein degradation machinery. Nat Struct Mol Biol 26, 946–954 (2019).

14. Fei, X., Bell, T.A., Barkow, S.R., Baker, T.A. & Sauer, R.T. Structural basis of ClpXP recognition and unfolding of ssrA-tagged substrates. Elife 9(2020).

15. Truscott, K.N., Lowth, B.R., Strack, P.R. & Dougan, D.A. Diverse functions of mitochondrial AAA+ proteins: protein activation, disaggregation, and degradation. Biochem Cell Biol 88, 97–108 (2010).

16. Lowth, B.R. et al. Substrate recognition and processing by a Walker B mutant of the human mitochondrial AAA+ protein CLPX. J Struct Biol 179, 193–201 (2012).

17. Kang, S.G. et al. Functional proteolytic complexes of the human mitochondrial ATP-dependent protease, hClpXP. J Biol Chem 277, 21095–102 (2002).

18. Liu, K., Ologbenla, A. & Houry, W.A. Dynamics of the ClpP serine protease: a model for self-compartmentalized proteases. Crit Rev Biochem Mol Biol 49, 400–12 (2014).

19. Kang, S.G., Maurizi, M.R., Thompson, M., Mueser, T. & Ahvazi, B. Crystallography and mutagenesis point to an essential role for the N-terminus of human mitochondrial ClpP. J Struct Biol 148, 338–52 (2004).

20. Brodie, E.J., Zhan, H., Saiyed, T., Truscott, K.N. & Dougan, D.A. Perrault syndrome type 3 caused by diverse molecular defects in CLPP. Sci Rep 8, 12862 (2018).

21. Wong, K.S. et al. Acyldepsipeptide Analogs Dysregulate Human Mitochondrial ClpP Protease Activity and Cause Apoptotic Cell Death. Cell Chem Biol 25, 1017–1030 e9 (2018).

22. Kang, S.G., Dimitrova, M.N., Ortega, J., Ginsburg, A. & Maurizi, M.R. Human mitochondrial ClpP is a stable heptamer that assembles into a tetradecamer in the presence of ClpX. J Biol Chem 280, 35424–32 (2005).

23. Wang, P. et al. Aberrant human ClpP activation disturbs mitochondrial proteome homeostasis to suppress pancreatic ductal adenocarcinoma. Cell Chem Biol 29, 1396–1408 e8 (2022).

24. Ishizawa, J. et al. Mitochondrial ClpP-Mediated Proteolysis Induces Selective Cancer Cell Lethality. Cancer Cell 35, 721–737 e9 (2019).

25. Puchades, C. et al. Unique Structural Features of the Mitochondrial AAA+ Protease AFG3L2 Reveal the Molecular Basis for Activity in Health and Disease. Mol Cell 75, 1073–1085 e6 (2019).

26. Puchades, C., Sandate, C.R. & Lander, G.C. The molecular principles governing the activity and functional diversity of AAA+ proteins. Nat Rev Mol Cell Biol 21, 43–58 (2020).

27. Shin, M. et al. Structures of the human LONP1 protease reveal regulatory steps involved in protease activation. Nat Commun 12, 3239 (2021).

28. Puchades, C. et al. Structure of the mitochondrial inner membrane AAA+ protease YME1 gives insight into substrate processing. Science 358(2017).

29. de la Pena, A.H., Goodall, E.A., Gates, S.N., Lander, G.C. & Martin, A. Substrate-engaged 26S proteasome structures reveal mechanisms for ATP-hydrolysis-driven translocation. Science 362(2018).

30. Shin, M.P. C.,; Asmita, A.; Puri, N.; Adjei, E.; Wiseman, R.L.; Karzai, A.W.; Lander G.C.;. Structural basis for distinct operational modes and protease activation in AAA protease Lon. Sci. Adv. 6(2020).

31. Ghanbarpour, A., Fei, X., Baker, T.A., Davis, J.H. & Sauer, R.T. The SspB adaptor drives structural changes in the AAA+ ClpXP protease during ssrA-tagged substrate delivery. Proc Natl Acad Sci U S A 120, e2219044120 (2023).

32. Ghanbarpour, A., Sauer, R.T. & Davis, J.H. A proteolytic AAA+ machine poised to unfold a protein substrate. Nature Communications 15 (2023).

33. Martin, A., Baker, T.A. & Sauer, R.T. Diverse pore loops of the AAA+ ClpX machine mediate unassisted and adaptor-dependent recognition of ssrA-tagged substrates. Mol Cell 29, 441–50 (2008).

34. Nagpal, J. et al. Molecular and structural insights into an asymmetric proteolytic complex (ClpP1P2) from Mycobacterium smegmatis. Sci Rep 9, 18019 (2019).

35. Goncalves, M.M. et al. Mechanism of allosteric activation in human mitochondrial ClpP protease. bioRxiv (2024).

36. Stahl, M. et al. Selective Activation of Human Caseinolytic Protease P (ClpP). Angew Chem Int Ed Engl 57, 14602–14607 (2018).

37. Malik, I.T. & Brotz-Oesterhelt, H. Conformational control of the bacterial Clp protease by natural product antibiotics. Nat Prod Rep 34, 815–831 (2017).

38. Felix, J., Weinhäupl, K., Chipot, C., Dehez, F., Hessel, A., Gauto, D.F., Morlot, C., Abian, O., Gutsche, I., Velazquez-Campoy, A., Schanda, P., Fraga, H.. Mechanism of the allosteric activation of the ClpP protease machinery by substrates and active-site inhibitors. Sci. Adv. 5(2019).

39. Mabanglo, M.F. et al. ClpP protease activation results from the reorganization of the electrostatic interaction networks at the entrance pores. Commun Biol 2, 410 (2019).

40. Baker, T.A. & Sauer, R.T. ClpXP, an ATP-powered unfolding and protein-degradation machine. Biochim Biophys Acta 1823, 15–28 (2012).

41. Strack, P.R. et al. Polymerase delta-interacting protein 38 (PDIP38) modulates the stability and activity of the mitochondrial AAA+ protease CLPXP. Commun Biol 3, 646 (2020).

42. Cottle, T., Joh, L., Posner, C., DeCosta, A. & Kardon, J.R. An adaptor for feedback regulation of heme biosynthesis by the mitochondrial protease CLPXP. bioRxiv (2024).

43. Burrows, P.C. et al. Functional roles of the pre-sensor I insertion sequence in an AAA+ bacterial enhancer binding protein. Mol Microbiol 73, 519–33 (2009).

44. Mabanglo, M.F. et al. Potent ClpP agonists with anticancer properties bind with improved structural complementarity and alter the mitochondrial N-terminome. Structure 31, 185–200 e10 (2023).

45. Tan, Y.Z. et al. Addressing preferred specimen orientation in single-particle cryo-EM through tilting. Nat Methods 14, 793–796 (2017).

46. Ghanbarpour, A. et al. A closed translocation channel in the substrate-free AAA+ ClpXP protease diminishes rogue degradation. Nat Commun 14, 7281 (2023).

47. Fux, A., Korotkov, V.S., Schneider, M., Antes, I. & Sieber, S.A. Chemical Cross-Linking Enables Drafting ClpXP Proximity Maps and Taking Snapshots of In Situ Interaction Networks. Cell Chem Biol 26, 48–59 e7 (2019).

48. Herzik, M.A.J. Setting up parallel illumination on the Talos Artica for high resolution data collection. Methods Mol Biol. 2215, 125–144 (2021).

49. Herzik, M., Wu, M. & Lander, G. Achieving better-than3-A-resolution by single-particle cryo-EM at 200-kev. Nat Methods 14, 1075–1078 (2017).

50. Punjani, A., Rubinstein, J.L., Fleet, D.J. & Brubaker, M.A. cryoSPARC: algorithms for rapid unsupervised cryo-EM structure determination. Nat Methods 14, 290–296 (2017).

51. Suloway, C. et al. Automated molecular microscopy: the new Leginon system. J Struct Biol 151, 41–60 (2005).

52. Lander, G.C. et al. Appion: an integrated, database-driven pipeline to facilitate EM image processing. J Struct Biol 166, 95–102 (2009).

53. Zheng, S.Q. et al. MotionCor2: anisotropic correction of beam-induced motion for improved cryo-electron microscopy. Nat Methods 14, 331–332 (2017).

54. Pettersen, E.F. et al. UCSF ChimeraX: Structure visualization for researchers, educators, and developers. Protein Sci 30, 70–82 (2021).

55. Emsley, P., Lohkamp, B., Scott, W.G. & Cowtan, K. Features and development of Coot. Acta Crystallogr D Biol Crystallogr 66, 486–501 (2010).

56. Adams, P.D. et al. PHENIX: a comprehensive Python-based system for macromolecular structure solution. Acta Crystallogr D Biol Crystallogr 66, 213–21 (2010).

57. Croll, T.I. ISOLDE: a physically realistic environment for model building into low-resolution electron-density maps. Acta Crystallogr D Struct Biol 74, 519–530 (2018).

